# The schizophrenia risk locus in SLC39A8 alters brain metal transport and plasma glycosylation

**DOI:** 10.1101/757088

**Authors:** Robert G. Mealer, Bruce G. Jenkins, Chia-Yen Chen, Mark J. Daly, Tian Ge, Sylvain Lehoux, Thorsten Marquardt, Christopher D. Palmer, Julien H. Park, Patrick J. Parsons, Robert Sackstein, Sarah E. Williams, Richard D. Cummings, Edward M. Scolnick, Jordan W. Smoller

## Abstract

A common missense variant in *SLC39A8* is convincingly associated with schizophrenia and several additional phenotypes. Homozygous loss-of-function mutations in *SLC39A8* result in undetectable serum manganese (Mn) and a Congenital Disorder of Glycosylation (CDG) due to the exquisite sensitivity of glycosyltransferases to Mn concentration. Here, we identified several Mn-related changes in human carriers of the common *SLC39A8* missense allele. Analysis of structural brain MRI scans showed a dose-dependent change in the ratio of T2w to T1w signal in several regions. Comprehensive trace element analysis confirmed a specific reduction of only serum Mn, and plasma protein N-glycome profiling revealed reduced complexity and branching. N-glycome profiling from two individuals with SLC39A8-CDG showed similar but more severe alterations in branching that improved with Mn supplementation, suggesting that the common variant exists on a spectrum of hypofunction with potential for reversibility. Characterizing the functional impact of this variant will enhance our understanding of schizophrenia pathogenesis and identify novel therapeutic targets and biomarkers of disease.

**Summary:** A common variant in the manganese transporter SLC39A8 is associated with numerous phenotypes including schizophrenia. Mealer *et. al.* presents an in-depth analysis of brain MRI and plasma glycomics in human carriers of the common variant, identifying several manganese-related changes with potential for diagnostic and therapeutic biomarker development.

## Introduction

GWAS identify hundreds of disease and phenotype-associated variants, providing researchers with new insights into the genetic architecture of complex human phenotypes including schizophrenia. However, translating genetic associations into a mechanistic understanding of disease and novel treatments remains a considerable challenge. One of the most pleotropic variants in the human genome is rs13107325 (C->T), a missense variant in *SLC39A8* that results in a substitution of threonine for alanine at position 391 (A391T) in exon 8. The minor allele (T) is associated with schizophrenia (Schizophrenia Working Group of the Psychiatric Genomics Consortium, 2014) as well as more than 30 unique traits including: changes in immune and growth traits (Zhang et al., 2018); increased HDL (Waterworth et al., 2010); increased risk of risk of inflammatory bowel disease and severe idiopathic scoliosis (Li et al., 2016a); and decreased serum manganese (Mn) (Ng et al., 2015), diastolic blood pressure (International Consortium for Blood Pressure Genome-Wide Association Studies et al., 2011), fluid intelligence (Hill et al., 2019), neurodevelopmental outcomes (Wahlberg et al., 2018), grey matter volume in multiple brain regions (Elliott et al., 2018), and Parkinson’s disease risk (Pickrell et al., 2016). The minor allele (T) of rs13107325 occurs at a frequency of ~8% in those of European descent, presumably from positive selection within this lineage following migration to colder climates (Li et al., 2016b).

SLC39A8 (aka ZIP8) is a transmembrane protein that cotransports divalent cations with bicarbonate; though it is capable of transporting Zn, Fe, Cu, Co, and Cd in cells, multiple studies suggest the primary physiologic role in humans is the transport of Mn (He et al., 2006; Wang et al., 2012; Choi et al., 2018). Manganese is an essential trace element for human health and affects neuronal function and development of dopaminergic neurons, although excess Mn is associated with disease (Chen et al., 2015; Horning et al., 2015; Kumar et al., 2014). In the brain, Mn is at highest concentrations in the striatum, and Mn toxicity, also known as *manganism*, is characterized by a Parkinsonian phenotype resulting from dysfunction of the nigrostriatal pathway (Horning et al., 2015). Manganese is a cofactor for many enzymes, including superoxide dismutase, glutamine synthetase, pyruvate carboxylase, and arginase, as well as many glycosyltransferases such as β(1,4)-galactosyltransferase (Ramakrishnan et al., 2006). A large number of Golgi glycosyltransferases, generally those containing a DxD metal-binding motif, uniquely require Mn as a cofactor (Breton et al., 2006; Chang et al., 2011). Glycosylation is a highly regulated, step-wise process of covalently attaching branched sugar polymers to proteins and lipids, and is known to affect nearly all biological pathways (Varki, 2017). Glycosylation plays a critical role in human disease, with over 125 known Mendelian conditions, termed congenital disorders of glycosylation (CDGs), associated with mutations in glycotransferases and related genes (Ng and Freeze, 2018). On proteins, glycans are most commonly attached to asparagine (N-linked) or serine/threonine (O-linked) residues. Disorders of N-glycosylation are divided into two distinct groups (type I and type II CDG) based on the observed transferrin glycosylation pattern, with type I indicating a defect in glycan assembly in the ER and type II suggestive of impaired modification of the side chain in the Golgi apparatus (Péanne et al., 2018; Abu Bakar et al., 2018).

Two case reports in 2015 demonstrated the importance of SLC39A8 in human disease, as individuals harboring rare homozygous mutations in SLC39A8 displayed a severe type II congenital disorder of glycosylation (SLC39A8-CDG) and near total absence of blood Mn (Park et al., 2015; Boycott et al., 2015). Other markers related to the Mn-dependent enzymes pyruvate carboxylase and glutamine synthetase were normal, suggesting a unique vulnerability of glycosylation enzymes to Mn concentration in SLC39A8-CDG. Importantly, a marker of impaired glycosylation and some clinical phenotypes such as seizure activity improved following supplementation with glycosylation precursors (galactose and uridine) or Mn (Park et al., 2015, 2018b). A recent study by Rader and colleagues showed that SLC39A8 regulated Mn homeostasis through uptake from bile in inducible- and liver-specific-knockout mice (Lin et al., 2017). Serum protein N-glycans analyzed by MALDI-TOF in these mice was suggestive of impaired N-glycosylation. Analysis of plasma N-glycans in human rs13107325 homozygous minor allele carriers (*n*=12) showed a slight but significant increase in one N-glycan precursor (monosialo-monogalacto-biantennary glycan, abbreviated A2G1S1), though a complete N-glycan analysis and information on the subjects was not reported (Lin et al., 2017).

The primary objective of our study was to measure Mn-related phenotypes with relevance to schizophrenia risk and brain function in human carriers of the *SLC39A8* missense variant. We identified a dose-dependent association of the schizophrenia risk allele with changes in the T2w/T1w ratio in several brain regions. Measurement of 23 serum trace elements confirmed the specific reduction of only serum Mn observed in other studies on this variant, and plasma protein N-glycosylation was altered in both heterozygous and homozygous carriers characterized by decreased branching and complexity of N-glycans. Analysis of SLC39A8-CDG plasma identified similar but more severe changes in protein N-glycosylation that were improved with Mn therapy, suggesting therapeutic intervention may be feasible in those carrying the *SLC39A8* risk allele.

## Results

### T2w/T1w ratios are changed in globus pallidus (GPi), lateral putamen (LPut), and substantia nigra (SN) in human A391T carriers

Previous studies have linked the rs13107325 minor allele (T) with brain MRI changes attributed to regional volumetric differences (Elliott et al., 2018; Luo et al., 2019). Given the paramagnetic properties of Mn and its known effect on MRI relaxation time (Pan et al., 2011; Malheiros et al., 2015; Lee et al., 2015), we hypothesized that the signal change resulted from changes in local concentrations of Mn and related ions. Decreased Mn would result in longer relaxation times of both T1- and T2- weighted images (T1w, T2w); however, longer T1 leads to lower signal intensity on T1w images, while longer T2 leads to increased signal on T2w images. We predicted that the ratio between the signal intensity of T2w and T1w images (T2w/T1w) would be a more sensitive parameter than either alone, with decreased Mn concentration in A391T carriers increasing in the T2w/T1w ratio. As UK Biobank images do not include T1 or T2 mapping as part of their protocol, comparing the ratio avoids problems with inter-subject normalization of T1w and T2w signal intensities due, for instance, to different coil loading factors and body size.

Brain MRI data were downloaded from the UK Biobank for 48 participants with homozygous minor allele genotype at rs13107325 (TT) along with 48 heterozygous minor (CT) and 48 homozygous major (CC) individuals matched on age, gender, smoking status, living area, BMI, and Townsend Deprivation Index as a proxy for socioeconomic status (Supp. Table 1). Due to problems with either the T1 or T2 images in some subjects, the final number of individuals with both scans included in the analysis were 47, 45, and 44 subjects for CC, CT, and TT genotypes, respectively. T2w/T1w ratios were compared between minor allele carriers (CT, TT) and controls (CC) using a t-test corrected for a false discovery rate of 5% on a pixel by pixel basis. In TT carriers, increased T2w/T1w ratios were observed in lateral putaminal (LPut) areas and diffusely through white matter (Fig. 1a). Contrary to our prediction, a decrease in the T2w/T1w ratio was observed in the globus pallidus interna (GPi) and substantia nigra (SN) of TT carriers, two areas of the brain with links to Mn toxicity and high levels of the divalent metal ion transporter DMT-1 (*SLC11A2)* (Erikson et al., 2004). CT carriers display changes in similar regions and in the same direction as TT carriers though with smaller effect size - only GPi and SN showed significant differences compared to CC genotype.

**Figure 1.**
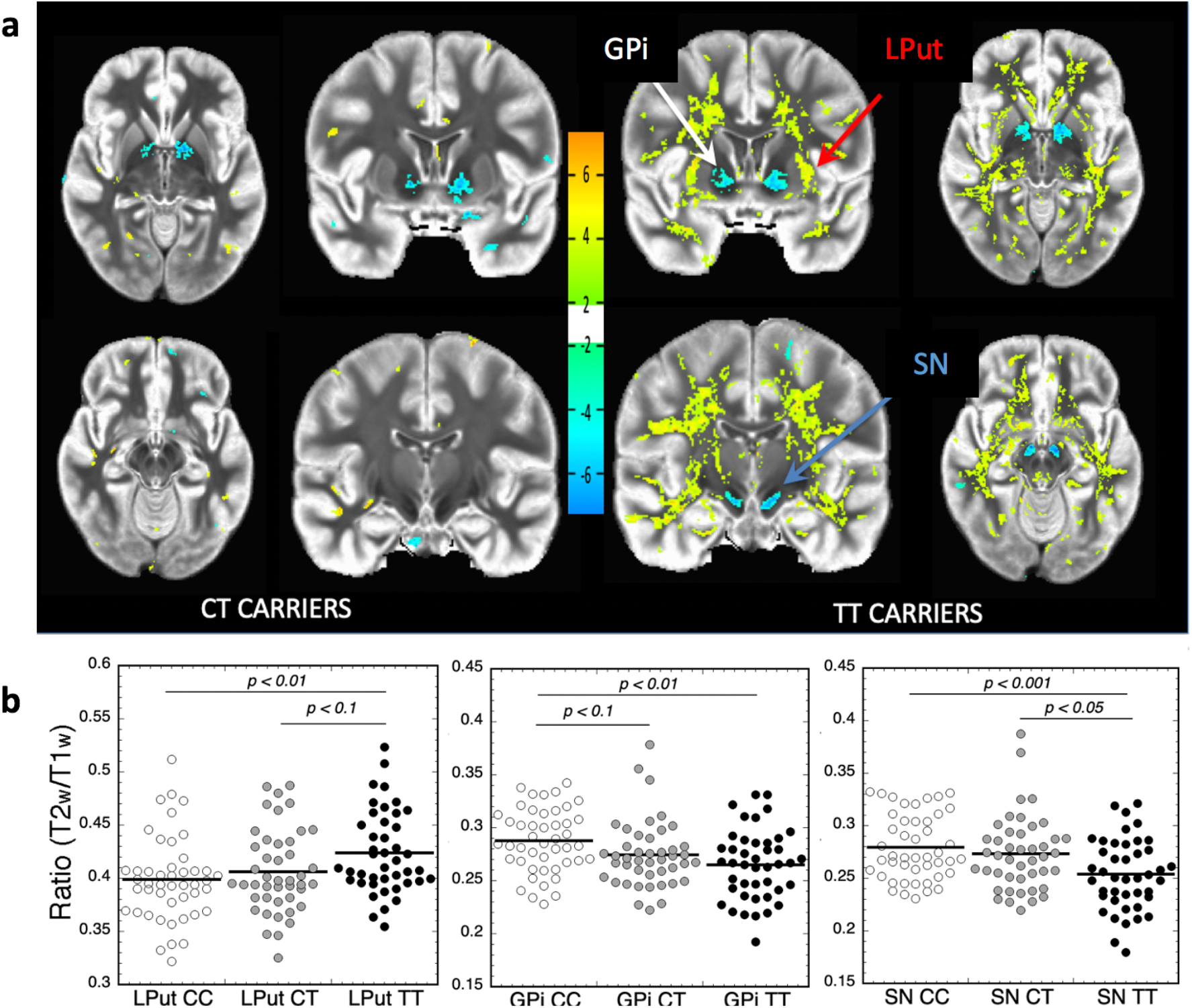
Dose-dependent effects of the A391T variant on regional T2w/T1w ratio. (**a**) Representative maps of the T2w/T1w ratio in CT and TT carriers overlaid on CC controls where ratio is either increased or decreased with a p-value of <0.05 on a pixel-by-pixel basis via student’s t-test. Heat map corresponds to the direction of change of the T2w/T1w ratio relative to the CC group, with yellow/orange representing an increase and blue representing a decrease. (**b**) Quantification of ROIs including globus pallidus (GPi), lateral putamen (LPut), and substantia nigra (SN) based on rs13107325 genotype, compared using post-hoc Dunnett’s test. Data points are shown for each individual, with the black horizontal line representing mean for each genotype. CC (white circles) n = 47, CT (gray circles) n = 45, TT (black circles) n = 44.

Dot plots for regions of interest (ROIs) are shown for GPi, SN, and LPut (Fig. 1b), with ANOVA analyses of the T2w/T1w ratios showing significant effects of genotype in GPi (F = 5.886; p = 0.0036; T2w/T1w ratio ± SD: CC = 0.279 ± 0.031, CT = 0.273 ± 0.035, TT = 0.253 ± 0.034; post-hoc Dunnett’s test: CC vs TT p = 0.0017; CC vs CT p = 0.09, and CT v TT p = 0.267), SN (F = 7.086; p = 0.0012; T2w/T1w ratio ± SD CC = 0.279±.031; CT = 0.275±.035; TT = 0.265±.033; Post-hoc Dunnett’s test: CC vs TT p = 0.0008; CC vs CT p = 0.617 CT v TT p = 0.013), and GPi (F = 4.72, p = 0.0105; T2w/T1w ratio ± SD: CC = 0.399±.039; CT = 0.406±.039; TT = 0.423±.038: Post-hoc Dunnett’s test: CC vs TT p = 0.0061; CC vs CT p = 0.566; CT v TT p = 0.071). These effects were driven by changes in both T1w signal and T2w signal because both were significantly different between CC vs TT genotypes, though the effect sizes were much larger for T2w signal in the GPi and SN. Additional analyses using MRI data to classify by genotype showed excellent separation of the CC and TT genotypes using only the T2w/T1w ratios of GPi, SN, and LPut (Supp. Fig. 1a, 1b). We note that use of raw T1w or T2w signal is not a useful comparator given the differences in BMI between the CC and TT genotypes. Regression analysis of all demographic data and T2w/T1w ratios found a significant correlation only between the LPut and BMI in TT carriers (Supp. Fig. 1c).

Though our sample size is smaller than prior studies (Elliott et al., 2018; Luo et al., 2019), a preliminary analysis using FreeSurfer software (http://surfer.nmr.mgh.harvard.edu/) showed no detectable difference in volume between genotypes in any brain region, suggesting that MRI signal changes in A391T carriers likely result from differences in the concentrations of paramagnetic ions such as Mn (data not shown).

### A391T is associated with a specific reduction of serum Mn

We next measured peripheral markers of Mn-related phenotypes in A391T carriers in samples matched for age and gender across genotypes available from the Partners Biobank (https://biobank.partners.org). Demographic characteristics of the sample are shown in Supp. Table 1. Serum concentrations of 23 trace elements were measured using Inductively Coupled Plasma-Mass Spectrometry (ICP-MS) and analyzed based on rs13107325 genotype. The method detection limit (MDL) for each element was determined on seven independent runs of the low-level quality control sample (Supp. Table 2). Measurements below the MDL are not distinguishable from background and have no quantitative confidence, limiting the reliability of detecting differences in concentration in this range. Of the trace elements measured, 16 had a mean across all samples above the MDL and were included in the analysis (As, Ba, Co, Cr, Cu, Cs, Hg, Mn, Mo, Pb, Sb, Se, Sn, U, V, and Zn), while 7 had a mean below the MDL and were excluded (Be, Cd, Ni, Pt, Te, Tl, and W) (Fig.2, Supp. Fig. 2)

**Figure 2.**
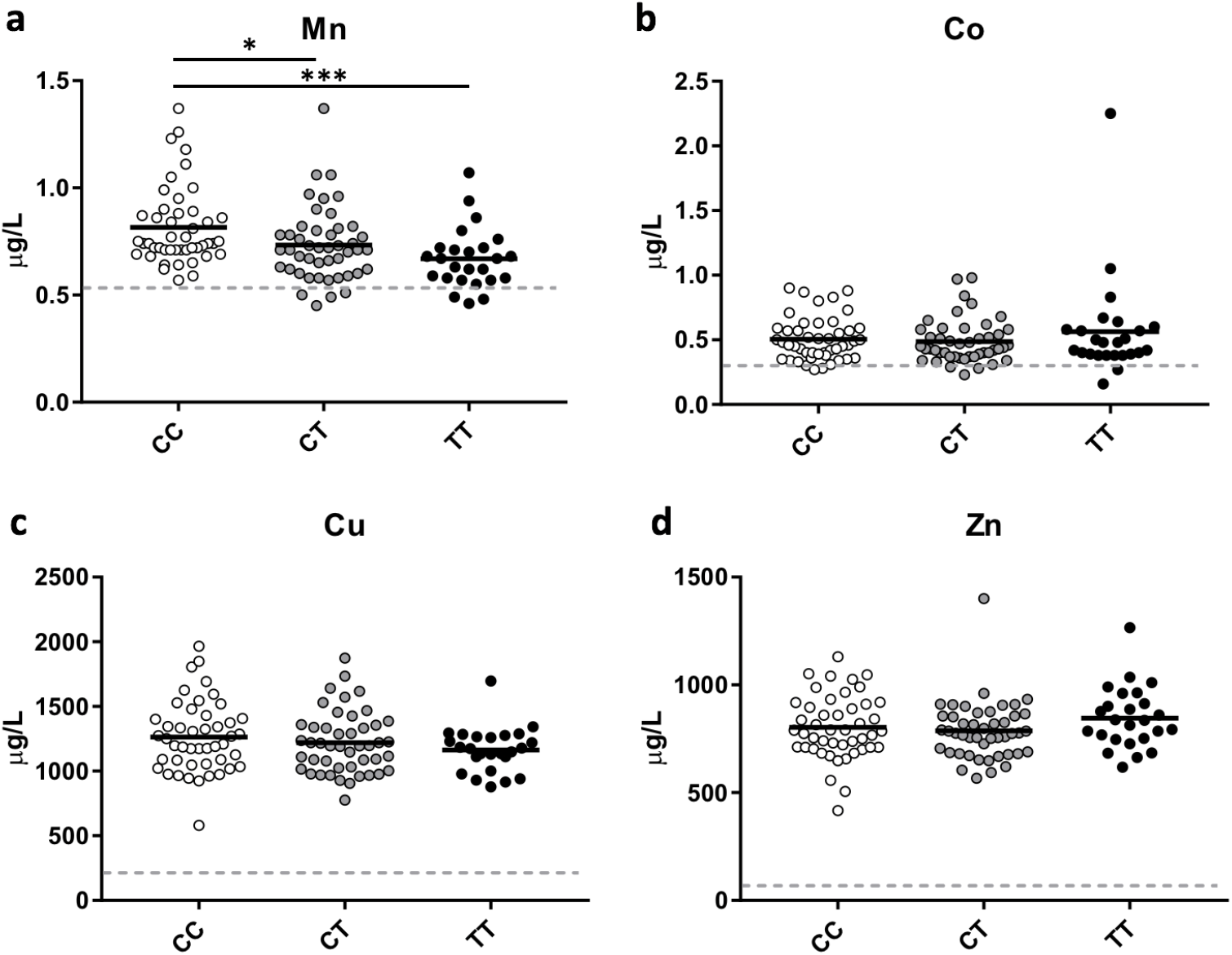
A391T results in a specific reduction of serum manganese in heterozygous and homozygous carriers. (**a**) Mn, (**b**) Co, (**c**) Cu, and (**d**) Zn are all trace elements previously shown to be transported by SLC39A8. Data points are shown for each individual, with the black horizontal line representing mean for each genotype. Method Detection Limit (MDL) for each trace element shown as grey dashed line on each graph. Genotypes compared using student’s t-test. CC (white circles) n = 46, CT (gray circles) n = 46, TT (black circles) n = 25. *p <0.05, **p <0.01, ***p <0.001.

ANOVA analyses each element showed that only Mn was statistically different between genotypes (F = 6.472, p = 0.0022, DF = 2). Heterozygous carriers of A391T showed a reduction of Mn concentration by 10% (CC 0.814 μg/L vs CT 0.732 μg/L, p = 0.027), whereas homozygous carriers had a reduction of 18% (CC 0.814 μg/L vs TT 0.669 μg/L, p = 0.0004) (Fig. 2a). The additional reduction of Mn in homozygous minor carriers suggests a dose-dependent effect of the mutation, though the difference between CT and TT groups fell short of statistical significance (CT vs TT p = 0.098). No difference was seen in the serum concentrations of other trace elements previously shown to be transported by SLC39A8 including Co, Cu, and Zn (Fig. 2b, 2c, and 2d, respectively).

Secondary sex-stratified analysis of genotype-related changes in serum Mn concentration showed a similar pattern but fell short of statistical significance in males (CT and TT) and TT females, likely due to the reduced sample size and the small effect size of the variant on Mn concentrations (Ng et al., 2015) (Supp. Fig. 3). There was no overall significant difference in serum Mn by sex, and linear regression showed no correlation between serum Mn concentration and age or BMI (Supp. Fig. 4).

### Analysis of plasma protein N-glycosylation showed reduced branching in CT and TT carriers

After confirming decreased serum Mn in A391T carriers from the Partners Biobank, we measured the plasma protein N-glycome based on rs13107325 genotype from the same individuals. N-glycans of plasma proteins were measured from 5 μL samples following peptide:N-glycosidase F (PNGaseF) cleavage and permethylation, and analyzed using MALDI-TOF MS based on standard protocols (Li et al., 2015a; Xia et al., 2013). Plasma was studied in lieu of serum given the abundance of literature on human plasma glycosylation (Clerc et al., 2016), and our pilot analysis of serum and plasma from the same donors found no substantial differences. Fifty-seven individual N-glycans were quantified after normalization for percent abundance within each sample. The overall N-glycome pattern, as illustrated by the 20 most abundant plasma N-glycans, was consistent with previous reports and overall similar between genotypes (Fig. 3a) (Clerc et al., 2016). Several individual glycans differed significantly based on genotype, and the direction of change was the same for the majority of individual N-glycans in CT and TT carriers (Supp. Table 3). A heat map of percent change (relative to CC) showed that larger glycans (m/z > 2851) consistently trend towards decreased abundance in CT and TT carriers (Fig. 3b).

**Figure 3.**
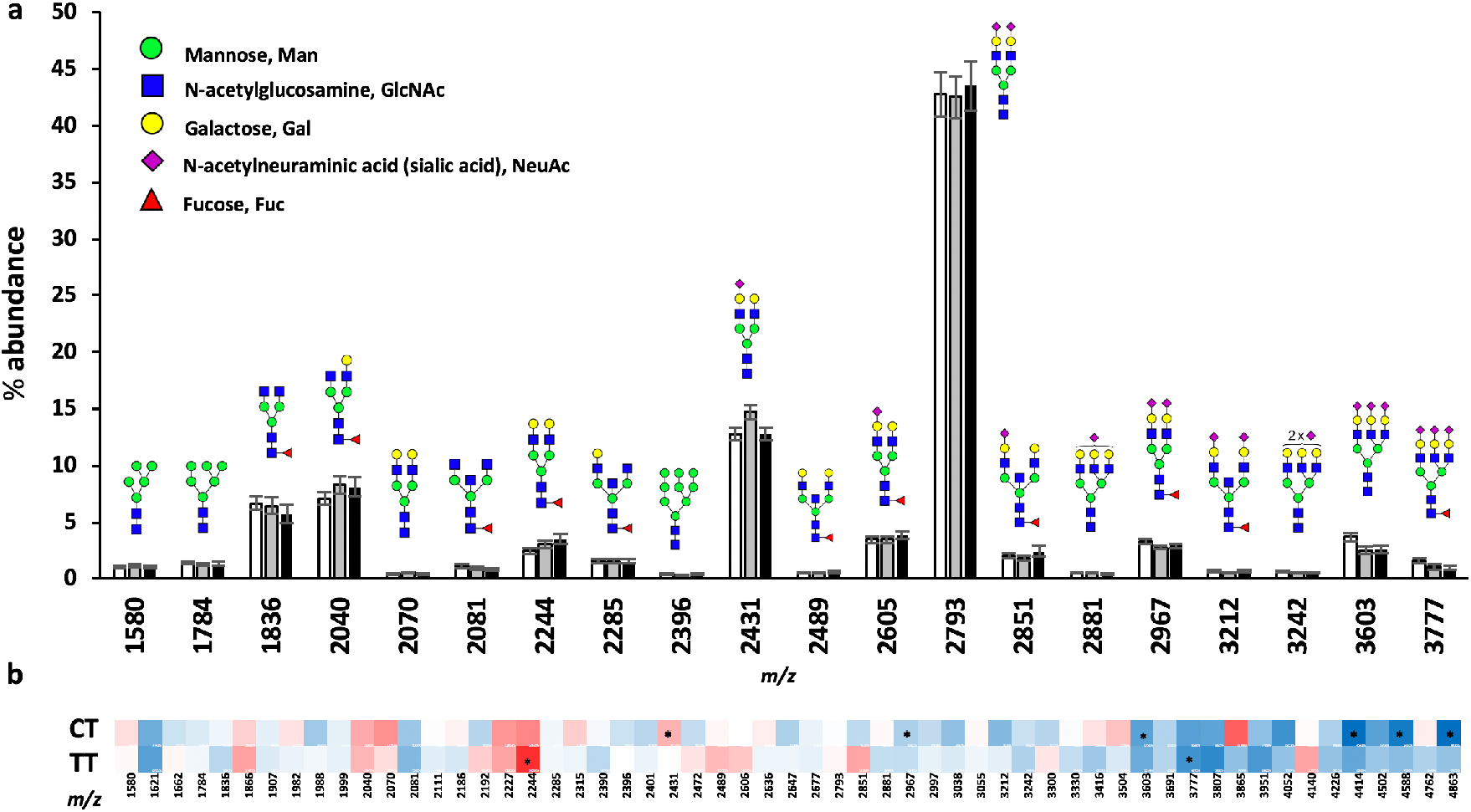
Analysis of plasma protein N-glycans based on rs13107325 genotype. (**a**) Summary data for the 20 most abundant plasma protein N-glycans sorted by rs13107325 genotype. Data presented as mean +/− standard error of the mean (SEM) for the percent (%) abundance of each N-glycan relative to the total N-glycan pool. Corresponding N-glycan structures are shown above each predicted m/z (mass/charge ratio) including a key for individual monosaccharide components of human N-glycans. (**b**) Heat map illustrating percent change of each individual glycan in CT and TT relative to CC; scaled from dark blue → white → bright red as −50.0% → 0% → +50.0%. Genotypes compared using student’s t-test. Individual N-glycans that are significantly different in CT and TT compared to CC are marked with an asterisk; *p <0.05. CC (white bars) n = 33, CT (gray bars) n = 31, TT (black bars) n = 25.

To determine whether specific enzymatic machinery is uniquely affected by genotype and Mn levels, glycans sharing structural similarities (such as branching, fucosylation, sialylation, etc.) were analyzed together, providing a more comprehensive evaluation than a single N-glycan. For example, bisection of N-glycans is performed by a single enzyme, MGAT3; analysis of all bisected N-glycans would be a more accurate readout of MGAT3 function than a single bisected N-glycan. Classification of each glycan by category is included in supplementary material (Supp. Table 4).

Branching of N-glycans, or antennarity, is defined as the number of N-acetylglucosamine (GlcNAc) linkages to core mannose (Man) residues, and is a proxy for the complexity of N-glycans (Varki et al., 2015). We observed reduced branching in CT and TT carriers compared to CC (Fig. 4; Table 1). In both CT and TT genotypes, there is a statistically significant increase in bi-antennary N-glycans (CC 88.0% vs CT 90.2%, p =0.0017; CC vs TT 90.1%, p =0.0082) and decrease in tri-antennary N-glycans (CC 6.77% vs CT 4.99%, p =0.0118; CC vs TT 4.91%, p =0.0126). Tetra-antennary glycans show a similar reduction in CT and TT carriers, though it did not reach statistical significance (CC 0.574% vs CT 0.420%, p =0.107; CC vs TT 0.457%, p =0.192). No change was observed in mono-antennary N-glycans and high-mannose structures (N-glycan precursors lacking antenna). Sex-stratified analysis revealed a greater effect on branching in male CT and TT carriers, though both sexes show a similar pattern (Fig. 5, Supp. Table 5).

**Table 1.**
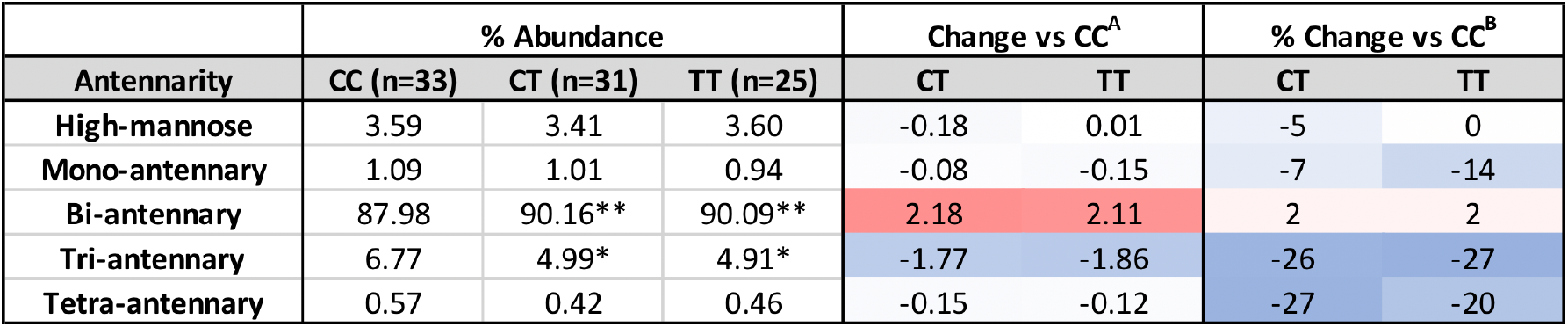
Plasma protein N-glycan branching based on rs13107325 genotype. Heat maps scale: dark blue -> white -> bright red representing −5.0 -> 0 -> +5.0 for absolute abundance^A^ change and −50.0% -> 0 -> +50.0% for relative change^B^. *p <0.05, **p <0.01 for % abundance of CT and TT vs CC genotype.

**Figure 4.**
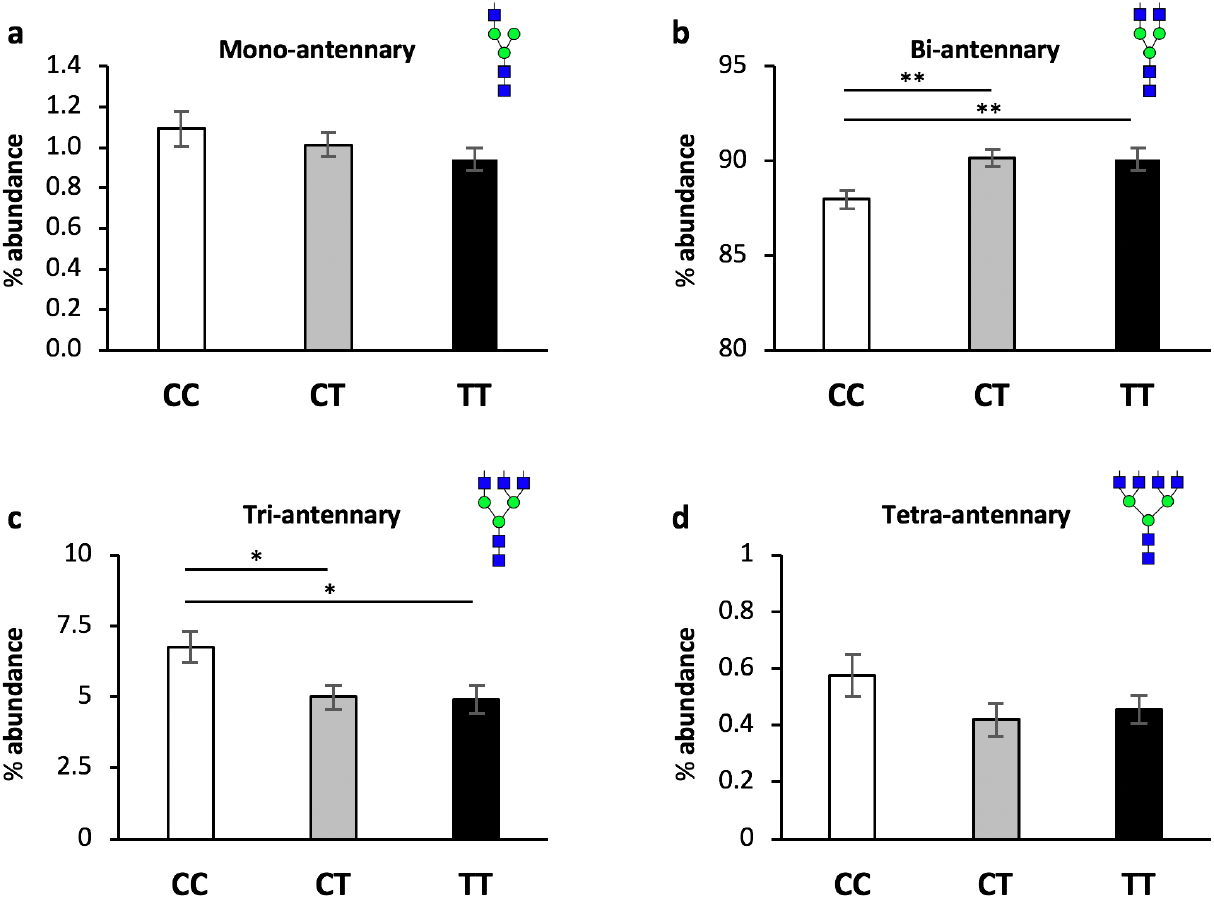
A391T carriers have reduced branching of plasma protein N-glycans. Data presented as mean +/− SEM for the % abundance of N-glycans with (**a**) one (mono-), (**b**) two (bi-), (**c**) three (tri-) or (**d**) four (tetra-) antennas, defined as the number of GlcNAc attachments to core Man residues (Supp. Table 3). Genotypes compared using student’s t-test. CC (white bars) n = 33, CT (gray bars) n = 31, TT (black bars) n = 25.

**Figure 5.**
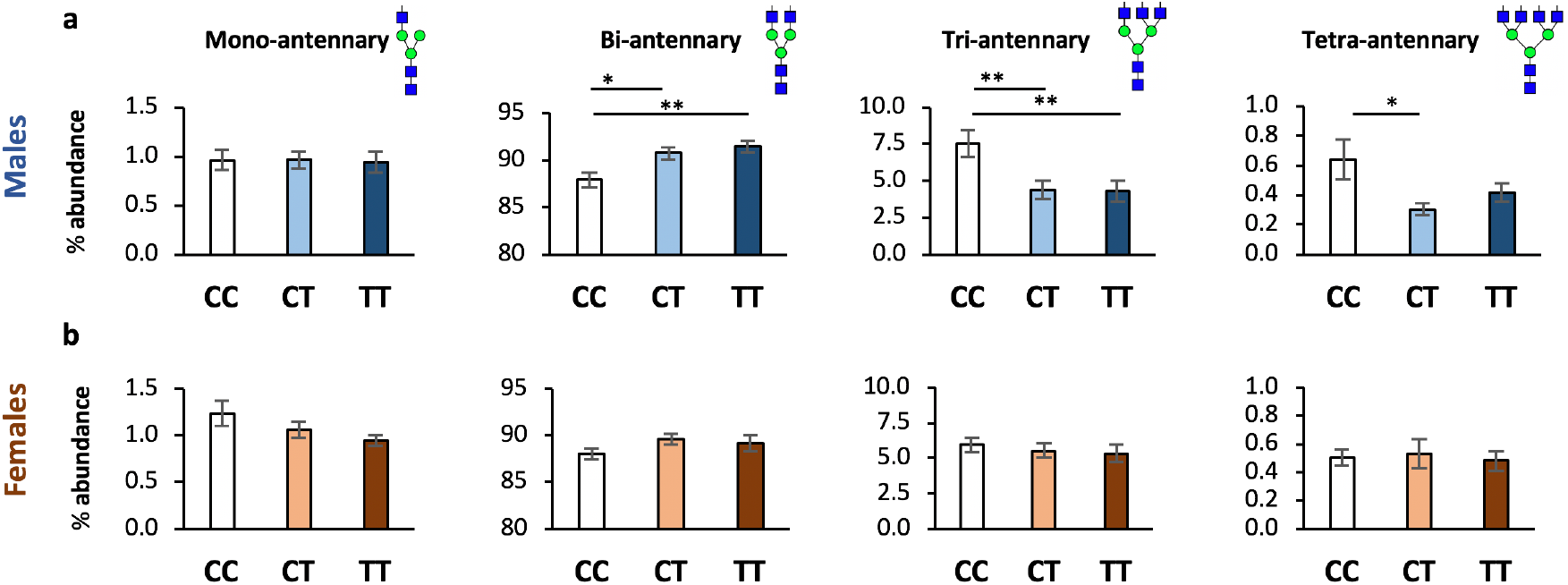
A391T has a larger effect on branching in male carriers. **a**, Males. **b**, Females. Data presented as mean +/− SEM for the percent abundance of N-glycans with one (mono-), two (bi-), three (tri-) or four (tetra-) antennas. Males: CC n = 17, CT n = 15, TT n = 10; Females: CC n = 16, CT n = 16, TT n = 12. *p-value <0.05, **p-value <0.01.

No significant change was observed in hybrid, bisecting, or core-fucosylated N-glycans (Supp. Table 6). Fucosylation of antenna, shown to be primarily tri- and tetra-antennary N-glycans (as opposed to core-fucosylation primarily on mono and bi-antennary structures) (Clerc et al., 2016), was reduced in both CT and TT carriers relative to CC, though only significantly in the TT group (CC 2.09% vs CT 01.56%, p =0.146; CC vs TT 1.43%, p =0.037). Analysis based on terminal monosaccharides showed no significant changes across any category, though there was a trend towards less sialylated species in CT and TT carriers (Supp. Table 6). Analysis stratified on the number of each residue showed a similar pattern of reduced abundance of larger and more complex structures in CT and TT carriers relative to CC. Finally, there was no change in the overall representation of each individual monosaccharide in the total protein N-glycan pool between genotypes, suggesting that differences in branching are not due to altered availability of the enzymatic substrate (UDP-Gal, UDP-GlcNAc, etc.) (Supp. Table 6).

### SLC39A8-CDG patients have increased precursor N-glycans and decreased complex N-glycans that are reversed following Mn supplementation

After completing glycome analysis of a common variant with a small effect, we sought to determine if more intolerant mutations in *SLC39A8* result in a similar but larger effect. Advances in next-generation DNA sequencing has resulted in an expansion of the known congenital disorders of glycosylation to well over one-hundred (Ng and Freeze, 2018). A pair of case studies in 2015 (Park et al., 2015; Boycott et al., 2015) and a recent report in 2017 (Riley et al., 2017) have identified multiple individuals with congenital disorders of glycosylation resulting from intolerant mutations in *SLC39A8* inherited in a recessive manner. The clinical phenotypes are dramatic and overlap in intellectual disability, seizures, brain structural abnormalities, low to undetectable Mn, and impaired transferrin glycosylation. Transferrin N-glycosylation is a common screening test for CDGs, though recent efforts have focused on performing mass spectrometry (MS) based methods given the more complete and sensitive nature of the test (Li et al., 2015a).

We performed plasma protein N-glycan profiling of two individuals with CDGs caused by homozygous SLC39A8 mutations before and after Mn supplementation. A full characterization of the clinical presentations as well as Mn supplementation protocol is described elsewhere (Park et al., 2018b). In brief: Subject A is an 8-month-old female with a more severe phenotype, harboring two mutations in highly conserved sites of *SLC39A8* (Gly38Arg, Ile340Asn), treated for ~1 year with Mn-sulfate after a cross-titration from galactose supplementation; Subject B is a 19-year-old female with a milder phenotype found to have 3 mutations in *SLC39A8* (Val33Met/Ser335Thr, Gly204Cys) treated with Mn-sulfate for ~1 year. The full plasma protein N-glycan profile and spectra for each individual pre- and post-Mn therapy is included in the supplementary material (Supp. Fig. 5, Supp. Table 7). Each sample was analyzed twice and produced similar results. We highlight specific glycans and groups of glycans which: 1) changed in the same direction in both subject A and subject B; 2) were similar to changes observed in other CDGs using MALDI-TOF; 3) changed with Mn treatment; and 4) showed parallel changes relative to those observed in A391T carriers. Similar to prior studies on CDGs, identification of a single or few individuals with a particular CDG limits direct and statistical comparison to meaningful matched control populations due to small sample size, age differences, and clinical variability. We include values for CC carriers using as comparators for our experimental protocols. As described in our methods, only serum from subject A pre-Mn supplementation was available and analyzed in lieu of plasma.

Relative abundance of A2G1S1, a monosialo-monogalacto bi-antennary N-glycan with permethylated m/z of 2227, is consistently elevated in plasma/serum across multiple CDGs (Li et al., 2015a), and was the only N-glycan reported as significantly elevated by Rader and colleagues in A391T homozygotes (Lin et al., 2017). Both subject A and subject B show increased A2G1S1 at baseline that decreased following Mn treatment (A 2.246% -> 0.652% with Mn; B 1.650% -> 0.622% with Mn; CC 0.439%) (Supp. Table 7). Two of the most abundant large N-glycans, A3G3S3 and A3FG3S3 (m/z 3603 and 3777, respectively) are consistently reduced in multiple CDGs(Li et al., 2015a). Both subject A and subject B have decreased A3G3S3 and subject A has lower A3FG3S3 at baseline; following Mn supplementation both A3G3S3 and A3FG3S3 increase in both individuals (A3G3S3: A 3.25% -> 8.03% with Mn; B 2.24% -> 5.71% with Mn; CC 3.68%), (A3FG3S3: A 0.662% -> 0.725% with Mn; B 3.81% -> 6.72% with Mn; CC 1.58%) (Fig. 6, Supp. Table 7).

**Figure 6.**
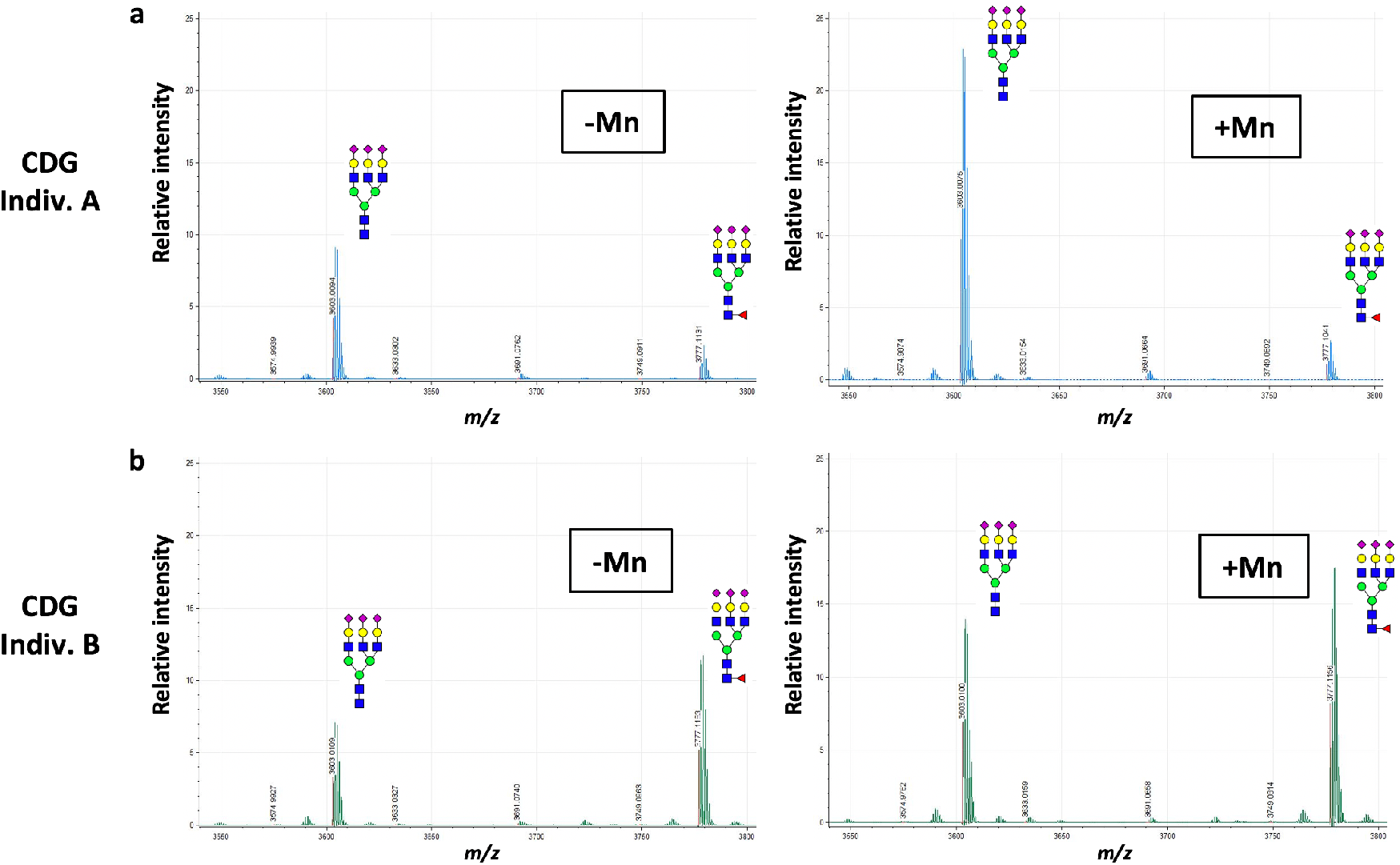
Mn supplementation increases large N-glycans in severe SLC39A8 mutation carriers with CDGs. Partial MALDI-TOF spectrum from plasma/serum of (**a**) subject A and (**b**) subject B pre- and post-Mn supplementation are shown. X-axis scaled for m/z of 3550-3800 kd and relative signal intensity on the Y-axis.

Grouped analysis of antennarity showed a decrease in mono- and bi-antennary structures and a dramatic increase in tri- and tetra-antennary structures following Mn treatment in both individuals (mono-antennary: A 1.49% -> 1.11% with Mn; B 1.06% -> 0.61% with Mn; CC 1.09%) (bi-antennary: A 88.8% -> 84.9% with Mn; B 87.7% -> 82.8% with Mn; CC 88.0%) (tri-antennary: A 5.48% -> 10.8% with Mn; B 8.22% -> 14.4% with Mn; CC 6.77%) (tetra-antennary: A 0.46% -> 0.78% with Mn; B 0.72% -> 0.89% with Mn; CC 0.57%) (Table 2, Fig. 7). High-mannose precursors were reduced following Mn treatment in both subjects (A 3.76% -> 2.39% with Mn; B 2.28% -> 1.31% with Mn; CC 3.59%). Bisecting N-glycans, synthesized only by MGAT3, which harbors a Mn-binding DxD motif, were markedly increased following Mn treatment (A 1.10% -> 3.90% with Mn; B 3.26% -> 4.35% with Mn; CC 7.83%) (Fig. 7). Analysis based on terminal monosaccharides showed more variable differences between the subjects without any clear trends in both subjects (Supp. Table 8). Analysis stratified by residue shows a similar pattern of increased abundance of more complex structures following Mn treatment, and the overall representation of each monosaccharide in the total N-glycan pool remained similar before and after Mn treatment aside from a 50% increase in fucose in subject A (Supp. Table 8). In summary, the MALDI-TOF N-glycan profiles of two individuals with SLC39A8-CDG showed reduced complexity of N-glycans, which was increased after one year of Mn supplementation.

**Table 2.**
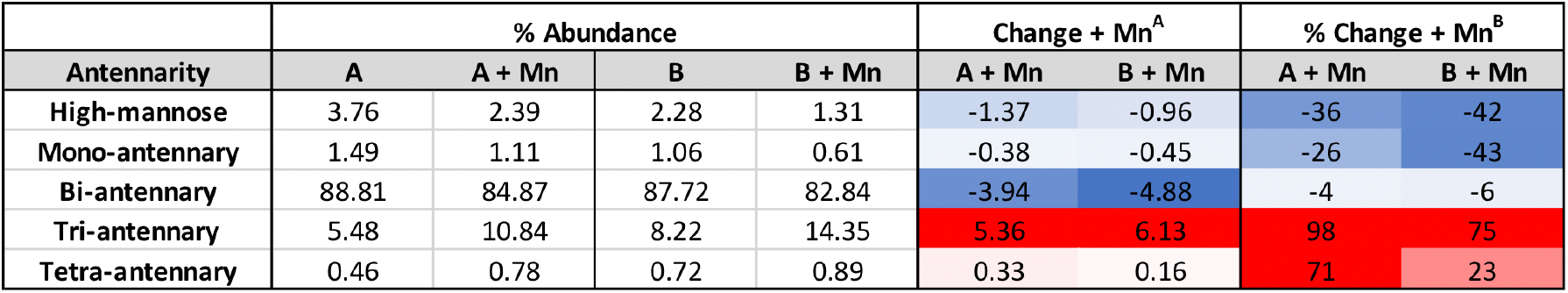
Plasma protein N-glycan branching following Mn supplementation in severe SLC39A8 mutation carriers. Heat maps scale: dark blue -> white -> bright red representing −5.0 -> 0 -> +5.0 for absolute abundance change^A^ and −50.0% -> 0 -> +50.0% for relative change^B^.

| Antennarity | % Abundance |  |  |  | Change + Mn<sup>A</sup> |  | % Change + Mn<sup>B</sup> |  |
| --- | --- | --- | --- | --- | --- | --- | --- | --- |
|  | A | A + Mn | B | B + Mn | A + Mn | B + Mn | A + Mn | B + Mn |
| High-mannose | 3.76 | 2.39 | 2.28 | 1.31 | -1.37 | -0.96 | -36 | -42 |
| Mono-antennary | 1.49 | 1.11 | 1.06 | 0.61 | -0.38 | -0.45 | -26 | -43 |
| Bi-antennary | 88.81 | 84.87 | 87.72 | 82.84 | -3.94 | -4.88 | -4 | -6 |
| Tri-antennary | 5.48 | 10.84 | 8.22 | 14.35 | 5.36 | 6.13 | 98 | 75 |
| Tetra-antennary | 0.46 | 0.78 | 0.72 | 0.89 | 0.33 | 0.16 | 71 | 23 |

**Figure 7.**
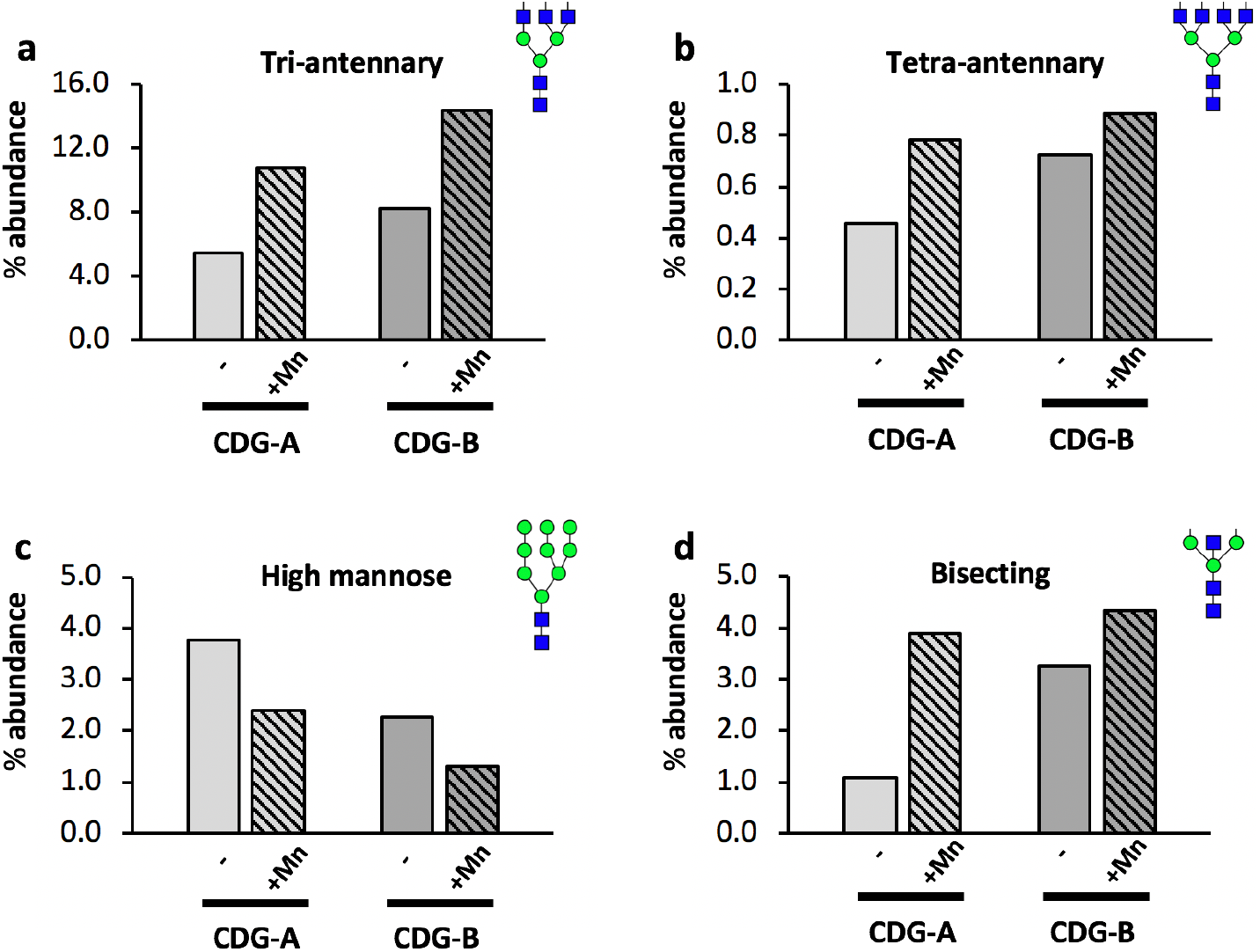
Plasma protein N-glycan changes in severe loss-of-function SLC39A8 mutation carriers after Mn treatment. Data presented as percent abundance of (**a**) tri-antennary, (**b**) tetra-antennary, (**c**) high-mannose, and (**d**) bisecting N-glycans before and after ~ 1 year of Mn supplementation in Subjects A and B with congenital disorders of glycosylation due to severe SLC39A8 homozygous mutations. Samples from each individual were replicated twice with similar results.

## Discussion

In our study, we identified several Mn-related changes in human carriers of a missense variant in *SCL39A8* associated with schizophrenia. In samples unaffected by schizophrenia, analysis of brain MRI data identified several regions of altered T2w/T1w signal based on rs13107325 genotype, consistent with changes in paramagnetic metal concentration. Both heterozygous and homozygous minor allele (A391T) carriers had a specific reduction of serum Mn and decreased branching of plasma protein N-glycans. Patients with SLC39A8-CDG showed a similar pattern of N-glycome changes that were reversed after a year of treatment with Mn, suggesting that glycome alterations and associated phenotypes in A391T carriers may be amenable to Mn supplementation. In particular, given the robust association of this variant with schizophrenia(Schizophrenia Working Group of the Psychiatric Genomics Consortium, 2014), our results raise the intriguing possibility that Mn supplementation might have therapeutic benefit for patients with this disorder.

The rs13107325 variant is associated with a diverse array of heritable traits and conditions. At the time of this manuscript preparation, the NHGRI-EBI GWAS Catalog (https://www.ebi.ac.uk/gwas/) lists 33 unique traits associated with rs13107325. Numerous associations relate to developmental processes, such as growth and immune traits, inflammatory bowel disease, scoliosis, intelligence, and schizophrenia, suggesting there may be critical windows of susceptibility to altered Mn concentration resulting from the missense variant.

We hypothesized that lower plasma Mn levels would be associated with lower brain Mn levels in rs13107325 minor allele carriers, leading to increased T1 and T2 relaxation times with higher T2w signal and lower T1w signal. We identified increased T2w/T1w ratios in putamen (primarily lateral putamen) and white matter tracts consistent with this hypothesis. In contrast, the GPi and SN showed decreased T2w/T1w ratios in both CT and TT carriers, suggestive of increased paramagnetic metal ion deposition. Mn transport in the brain is complex, regulated by numerous transporters and pathways with significant overlap between iron (Fe) and Mn homeostasis (Chen et al., 2015; Horning et al., 2015; Erikson et al., 2004; Aschner and Aschner, 1991; Chen et al., 1989). Both increased and decreased Fe lead to increased Mn transport into the brain (Fitsanakis et al., 2010, 2011). We suspect that low levels of Mn lead to increased uptake of Fe in some regions, particularly those with high affinity for divalent cations such as the GPi and SN. The GPi has high levels of DMT-1 expression and the highest rate of Mn deposition in welders exposed to Mn (Lee et al., 2015). Studies in primates have demonstrated that the pallidum is uniquely susceptible to Mn accumulation (Dorman et al., 2006; Sung et al., 2007), and manganese toxicity results in a Parkinsonian phenotype (16,17). Interestingly, A391T is protective against Parkinson disorder, presumably from reduced uptake of Mn in to the pallidum or SN (Pickrell et al., 2016). Two recent studies associated A391T with brain MRI signal changes attributed to volumetric differences including increased gray matter in the caudate, putamen and cerebellum (Elliott et al., 2018; Luo et al., 2019). MRI signal is affected by any factor influencing the magnetic field including Fe and Mn concentrations (Duyn, 2013). Voxel-based morphometry (VBM) detects intensity changes between brain regions and is often interpreted as changes in grey matter density and volume. In contrast to VBM techniques, our preliminary studies measured volume using FreeSurfer and found no volumetric difference in any brain region based on rs13107325 genotype. We conclude that the direction of the T2w/T1w signal change in A391T carriers resulted from regional changes in Mn, Fe, or both.

Our study confirms that the A391T missense mutation selectively lowers serum Mn levels in both heterozygous and homozygous carriers. No differences were detected in any of the other trace elements transported through *SLC39A8* (Zn, Co, and Cu), though effects on Cd could not be assessed as levels were below our method of detection limit. Fe was not measured in our study, though GWAS on Fe levels and Fe-related traits have not identified any associations with the A391T missense variant, and two recent studies show no difference in Fe concentration in A391T carriers, suggesting that serum Fe is not affected (Haller et al., 2018; Wahlberg et al., 2018)

Multiple studies demonstrate the importance of *SLC39A8* to health and disease. Hypomorphic mice expressing 10-15% of basal SLC39A8 show severe growth stunting, dysmorphogenesis, and anemia (Gálvez-Peralta et al., 2012, 8; Chen et al., 2018, 8). SLC39A8 is required during cardiac development and regulates zinc transport in endothelium (Lin et al., 2018). Recent studies also highlight a role for the *SLC39A8* common variant in regulation of the gut microbiome and metal homeostasis in Crohn’s disease (Li et al., 2016a; Collij et al., 2019; Melia et al., 2019). Two case series from 2015 identified individuals with severe mutations in *SLC39A8* presenting with a constellation of severe symptoms including intellectual disability, developmental delay, cerebellar atrophy, growth abnormalities, and seizures (Park et al., 2015; Boycott et al., 2015). A more recent report identified a pair of siblings presenting with a Leigh-like syndrome of intellectual disability, dystonia, seizures, cortical atrophy and basal ganglia T2 hyperintensities (Riley et al., 2017). Importantly, these studies demonstrate that the glycosylation deficits and some of the clinical phenotypes can be improved by oral supplementation of galactose and uridine to yield UDP-Gal, the key enzymatic substrate for β(1,4)-galactosyltransferase, as well as the attendant obligatory cofactor for this activity, Mn (Park et al., 2015; Riley et al., 2017; Park et al., 2018b).

Human plasma protein N-glycosylation is an extensive area of research, with detailed descriptions of the abundance of each glycan, identification of proteins harboring each glycan, and how these glycans affect protein function (Clerc et al., 2016). The plasma N-glycome is increasingly explored as a potential biomarker in a variety of settings including depression (Boeck et al., 2018; Park et al., 2018a; Yamagata et al., 2018), pregnancy (Jansen et al., 2016), IBD (Clerc et al., 2018), Down syndrome (Borelli et al., 2015), inflammation and metabolic health (Reiding et al., 2017), and post-surgical changes (Gudelj et al., 2016). Given the repeated association of *SLC39A8* with glycosylation defects in prior studies (Park et al., 2015; Boycott et al., 2015; Park et al., 2018b; Riley et al., 2017, 8; Lin et al., 2017), and the exquisite sensitivity of certain glycosyltransferases for Mn as an irreplaceable co-factor, we focused our study on glycosylation changes in A391T carriers.

Plasma protein N-glycome changes were similar in individuals carrying one or two copies of the hypo-functioning allele, suggesting a dominant effect. The most notable findings in individuals carrying the missense mutation were reduced branching and decreased complexity of large N-glycans. Rader and colleagues describe a small but significant increase in the abundance of the monosialo-monogalacto-biantennary precursor glycan A2G1S1 (*m/z* 2227) in a group of A391T homozygotes carriers (TT) but did not report the abundance of other plasma N-glycans (Lin et al., 2017). A2G1S1 is commonly increased in CDGs and suggestive of decreased β(1,4)-galactosyltransferase activity (Li et al., 2015b). We observed a ~20% increased abundance of A2G1S1 in both CT and TT carriers, though it did not reach statistical significance. We hypothesize that reduced Mn availability in mutation carriers results in a modest but broad reduction of glycosyltransferase activity in glycosyltransferases containing DxD domains including β(1,4)-galactosyltransferase and the numerous N-acetylglucosaminyltransferases (MGATs) that control N-glycan branching. In addition, the activity of some glycosyltransferases lacking a classic DxD domain are also affected by variations in Mn concentration including sialyltransferases and may be affected (Audry et al., 2011).

We report the first plasma N-glycome analysis using MALDI-TOF MS of two individuals with severe *SLC39A8* mutations causing CDGs before and after Mn supplementation(Park et al., 2015, 2018b). In plasma from these patients, we observed an elevation of the precursor A2G1S1 (*m/z* 2227) and a reduction of larger, more complex glycans including A3G3S3 and A3FG3S3 (*m/z* 3603 and 3777, respectively), similar to what has been reported in other type-II CDGs (Li et al., 2015a). The abundance of these three individual glycans normalized following Mn supplementation, and grouped analysis showed a dramatic increase in branched N-glycans and reduced high-mannose precursors following treatment. These changes parallel the reduced complexity of N-glycans in A391T carriers and suggest that such changes could be targeted with Mn supplementation. When and how to safely and effectively administer such a treatment remains to be determined. However, tracking of the branching of N-glycans could be a useful biomarker of treatment response and dose titration in CDGs and conditions associated with the A391T variant such as schizophrenia.

Glycosyltransferases associated with common, complex diseases tend to have specific tissue-specific expression profiles, isoenzymes with redundant activity, and function across multiple pathways, whereas glycosyltransferases associated with CDGs tend to have diffuse expression, lack redundant isoenzymes, and function within specific glycosylation pathways (Joshi et al., 2018). A minority of glycosyltransferases are associated with both common diseases and CDGs, similar to what is observed with *SLC39A8.* GWAS have identified loci influencing the glycosylation of specific plasma proteins such as IgG as well as the total glycome (Lauc et al., 2013; Wahl et al., 2018; Sharapov et al., 2019). These studies do not report an association of rs13107325 with plasma and IgG N-glycosylation patterns and primarily identify SNPs near glycosyltransferase genes. This may result from differences in methodologies employed between studies (MALDI-TOF vs UPLC (Sharapov et al., 2019) and LC-ESI-MS (Wahl et al., 2018)), minor allele frequency between cohorts, power/sample size, and effect size; rs13107325 may not be a major regulator of total plasma N-glycosylation relative to all genetic variation in glycosylation-related genes. Large, branched N-glycans are not generally found on IgG (Clerc et al., 2016), thus it is less likely changes associated with rs13107325 would be identified in such studies. Though MALDI-TOF MS is a standard tool in the study of glycosylation disorders, the semi-quantitative of this assay only allows determination of abundance changes within each sample after normalization. In addition to quantitative assays, future studies of SLC39A8-A391T should include analysis of protein O-glycans and glycolipids, as well as disease-relevant systems including primary tissue, cell lines, and murine models to determine how this variant elicits such pleotropic effects.

In summary, we have demonstrated that a common missense variant in *SLC39A8* is associated with multiple Mn-related phenotypes in both heterozygous and homozygous carriers, including reduced serum Mn levels, and decreased branching and complexity of plasma N-glycans. In addition, we identify parallel changes in the plasma N-glycome of SLC39A8-CDG patients that are reversed following Mn supplementation. Although the effect size of the common variant in *SLC39A8* on glycosylation is relatively modest, translation of validated genetic variants to functional biologic pathways can provide critical insight for a disease.

For example, common variants in *HMGCR* (gene for 3-hydroxy-3-methylglutaryl-CoA reductase) result in a modest effect on LDL levels despite this protein being the target for the majority of lipid-lowering medications (Kathiresan et al., 2009, 2008; Teslovich et al., 2010; Aulchenko et al., 2009), and a common variant in *DRD2*, the gene encoding the dopamine receptor D2 and the site of action for anti-psychotic medications (van Rossum, 1966; Enna et al., 1976), results in only a small increase in the risk of schizophrenia (Schizophrenia Working Group of the Psychiatric Genomics Consortium, 2014). Our observations provide mechanistic insights across a broad range of conditions including schizophrenia and may have implications for the development of novel diagnostic and therapeutic biomarkers.

## Methods

**MRI data** (T1 and T2 images) were downloaded from the UK Biobank 2018 release of ~15,000 participants’ imaging data (https://www.ukbiobank.ac.uk). The UK Biobank imaging protocol has been described previously (Elliott et al., 2018). Equal numbers (n = 48) of individuals with each of the three rs13107325 genotypes and brain imaging data were identified after matching for age, sex, smoking status, living area, body mass index (BMI) and Townsend Deprivation Index as a proxy for socioeconomic status. Ratio images were created by dividing the T2-weighted images (T2w) by the T1-weighted (T1w) images. Ratio images in TT and CT carriers were compared to CC subjects on a pixel-by-pixel basis using a t-test corrected for a false discovery rate of 5% using standard tools in the AFNI image analysis program (https://afni.nimh.nih.gov). Regions of interest (ROI), including the substantia nigra (SN), globus pallidus interna (GPi) and lateral putamen (LPut), were drawn after averaging all the images from the CC cohort, and then propagated to each individual ratio image. Values determined over the whole ROI were then compared using ANOVA analyses as described in Statistical Analysis, below.

**Serum and plasma samples** were obtained from the Partners Biobank, a biorepository containing serum, plasma, DNA, and buffy coats from 80,000 participants linked to the electronic health records (EHRs) and consented for broad-based research including 20,000 participants with genome-wide genotype data (https://biobank.partners.org). De-identified samples of serum and plasma were selected from 25 participants with the homozygous minor TT genotype independent of clinical phenotype, along with age- and sex-matched samples of the CC and CT genotype (46 each). Samples from each individual were analyzed by ICP-MS for trace elements concentration. Plasma N-glycomics analysis were processed in batches of 12, with each batch including at least 3 samples from each genotype. This resulted in the total number of analyzed samples reaching 33, 31, and 25 for CC, CT, and TT, respectively, when all TT samples had been analyzed. Serum and plasma samples were coded and blinded for all experiments with genotype revealed only for analysis. Demographic characteristics based on genotype are shown in Supplementary Table 1. Plasma and serum were provided by Drs. Park and Marquardt from two individuals with a CDG associated with mutations in *SLC39A8*, as described previously (Park et al., 2015, 2018b). Of note, the plasma sample provided for Subject A pre-Mn supplementation did not produce an interpretable N-glycan profile despite two repeat analyses as the signal intensity was too low, presumably due to a problem with storage or transfer of the sample. An available serum sample of Subject A pre-Mn supplementation was analyzed in lieu of plasma and produced a reliable and replicable N-glycan profile. Given the limited quantity of samples available from these rare-disease cases, and our observation that serum and plasma have similar N-glycan profiles as described above, the serum sample from Subject A was included in our report.

**Inductively Coupled Plasma-Mass Spectrometry (ICP-MS) Trace Element Analysis** was performed at the Wadsworth Center, New York State Department of Health (Albany, New York). For each of the 23 trace elements measured, the MDLs were determined on seven independent runs and appropriate, multi-level quality control (QC) samples included with each run. The QC data are shown in Supplementary Table 2 and are represented on each graph by a hashed line in Figure 2 and Supplemental Figure 2. For each sample, 200 μL of serum was diluted with a reagent containing appropriate internal standards for the analysis. Because serum samples in the Partners Biobank were obtained in BD-red top vacutainer tubes and aliquoted into cryovials for storage (as opposed to the BD-Royal Blue trace elements tubes), we performed a pilot study of four serum samples drawn simultaneously into red-top and royal blue top tubes, and then aliquoted into either cryovials or metal-free specimen vials. There was no difference in the measured Mn content between these two sample tubes, which suggests that archived sera in the Biobank is suitable for trace Mn measurements.

**Purification of plasma protein N-glycans** was performed using standard protocols consistent with prior studies on CDGs (Li et al., 2015a) and are available through The National Center for Functional Glycomics website (www.ncfg.hms.harvard.edu). In brief, 5 μL of plasma was lyophilized and resuspended in 20 μL 1X Rapid PNGaseF buffer (NEB #P0710S) and incubated for 15 minutes at 70°C to denature proteins. After cooling to room temperature, 1 μL of Rapid PNGaseF (NEB #P0710S) was added and incubated at 50°C for 1 hour to cleave N-glycans from proteins. PNGaseF treated samples were resuspended in 100 μl of 5% acetic acid and added to a C18 Sep-Pak (50 mg) column (Waters, #WAT054955) preconditioned with one column volume each of methanol, 5% acetic acid, 1-propanol, and 5% acetic acid. The reaction tube was washed with another 100 μL of 5% acetic acid and added to the C-18 column, followed by 1 mL of 5% acetic acid, and the entire flow-through was collected in a microcentrifuge tube (~1.2 mL). Samples were placed in a speed vacuum for two hours to reduce the volume to ~300 μL, covered in parafilm and lyophilized overnight.

**N-glycan permethylation** was performed using a fresh slurry of NaOH/DMSO daily. Seven pellets of NaOH (Sigma-Aldrich, #S8045) were dissolved in four glass pipettes volumes (~3 ml) of DMSO (Sigma-Aldrich, #D8418) and ground using a clean/dry mortar and pestle. 200 μL of the NaOH/DMSO slurry was added to the lyophilized N-glycans in addition to 100μL iodomethane (Sigma-Aldrich, #289566) and placed in on a vortex shaker for 20 minutes at room temperature with a microtube cap to prevent the lid from opening due to increased gas pressure. After the mixture became white, semi-solid and chalky, 200 μL ddH_2_O was added to stop the reaction and dissolve the sample. 200 μL chloroform and an additional 400 μL ddH_2_O were added for chloroform extraction and vortexed followed by brief centrifugation. The aqueous phase was discarded, and the chloroform fraction was washed three additional times with 800 μL ddH_2_O. Chloroform was then evaporated by 20 minutes in a speed vacuum. Permethylated N-glycans were resuspended in 200 μL of 50% methanol and added to a C18 Sep-Pak (50 mg) column preconditioned with one column volume each of methanol, ddH_2_O, acetonitrile, and ddH_2_O. The reaction tube was washed with 1mL 10% acetonitrile and added to the column, followed by an additional 2 ml wash of 10% acetonitrile. Columns were placed in a 15 mL glass tube, and permethylated N-glycans were eluted with 3 mL 50% acetonitrile. The eluted fraction was placed in a speed vacuum for one hour to remove the acetonitrile, covered in parafilm and lyophilized overnight.

**MALDI-TOF analysis of purified glycans** was performed on permethylated N-glycans resuspended in 25 μL of 75% methanol and spotted in a 1:1 ratio with DHB matrix on a metal 384 spot. Spectra from the samples were obtained in a Bruker MALDI-TOF instrument using FlexControl Software in the Reflection Positive mode with a mass/charge (m/z) range of 1,500-5,000 kD. Twenty independent captures (representing 1,000 shots each) were obtained from each sample and averaged to create the final spectra file and exported in .msd format for analysis.

**N-Glycan analysis.** 57 plasma N-glycans of known structure corresponding to the correct isotopic mass were annotated in each spectra using mMass software (Strohalm et al., 2008). The relative abundance of each N-glycan was calculated as the signal intensity for each peak divided by the signal intensity for all 57 measured N-glycans within a spectrum. N-glycans were grouped into different categories based on shared components such as monosaccharide composition, antennarity, or class based on deductive reasoning and prior MS/MS data where available (Clerc et al., 2016) (Supp. Table 4). Absolute change and relative change compared either to CC genotype or pre-Mn supplementation is shown. Heat maps are scaled from dark blue → white → bright red representing (−5.0 → 0 → +5.0) for absolute abundance change and (−50.0% → 0 → +50.0%) for relative change. The contribution of each monosaccharide was determined by taking the percentage of each monosaccharide in a N-glycan multiplied by the abundance of the glycan, and then summated for the five monosaccharides present in human plasma N-glycans.

**Statistical Analysis.** Brain MRI data of T2w/T1w ratios from TT and CT carriers were compared to CC subjects on a pixel by pixel basis using a t-test corrected for a false discovery rate of 5% using AFNI Image Analysis Tools (https://afni.nimh.nih.gov) and StatistiXL Version 2 Software. Regions of interest were compared as ratios of T2w/T1w signal intensities using a one-way ANOVA followed by a post-hoc comparison using a Dunnett’s test, with the CC as a control (or TT for comparisons between TT and CT). Metal data was analyzed using GraphPad Prism Version 7 and included an initial ANOVA analysis (degrees of freedom, DF = 2) followed by individual unpaired t-tests assuming unequal variance between each genotype (CC vs CT, CC vs TT, CT vs TT), followed by linear regression between Mn concentration, age, and BMI. Glycosylation data was analyzed using Microsoft Excel Version 16.27. The abundance of individual glycans and glycan classes were compared between genotypes using unpaired t-tests assuming unequal variance between genotypes, with significance thresholds applied at p *<0.05, **<0.01, and ***<0.001.

**Study Approval** was obtained from the Massachusetts General Hospital/Partners Human Research Committee IRB. Informed consent and approval for Mn supplementation in subjects with SLC39A8-CDG were obtained and described in prior publications (Park et al., 2015, 2018b).

**Supplemental Material** is available online, including: linear discrimination analysis of MRI data by genotype; complete trace element analyses and method detection limits for each element; description of glycan classifications used in this study; additional analysis of trace element and glycomics data based on gender, age, and BMI; and summary results of N-glycomics from based on rs13107325 genotype and in SLC39A8-CDGs.

## Author contributions

RGM directed and designed the project, performed all glycomics experiments and analysis, coordinated collaborations, and wrote the manuscript

BGJ performed the analysis of MRI data and helped write this portion of the manuscript MJD helped conceptualize the study based on GWAS results

TG collected and curated the MRI data from the UK Biobank

SL assisted with experimental methods of N-glycosylation and advised on analysis of glycans TM and JHP provided plasma/serum samples of subjects with SLC39A8-CDG

CDP performed trace element analysis of serum samples by ICP-MS

PJP supervised CDP and oversaw the ICP-MS trace element analyses, planned pilot studies for Mn contamination, and helped write the sections on trace element analysis

RS helped conceptualize the study based on GWAS results and link to glycosylation SEW performed statistical analyses and assisted with data preparation

RDC oversaw N-glycosylation analysis, helped conceptualize the study based on GWAS results and link to glycosylation, and supervised RGM on glycosylation aspects of the project

EMS helped conceptualize the study based on GWAS results and link to glycosylation, helped design the study, coordinated collaborations, and supervised RGM on all aspects of the project JWS oversaw the project, helped conceptualize study design, coordinated collaborations, and supervised RGM on all aspects of the project

All authors provided edits and feedback during preparation of the manuscript

## Acknowledgements

We would like to thank the Partners Biobank employees and participants for their work and contribution to this valuable research resource utilized in our study. This work was supported by a foundation grant from the Stanley Center for Psychiatric Research at the Broad Institute of Harvard/MIT (awarded to RGM). JWS is a Tepper Family MGH Research Scholar. This research has been conducted using the UK Biobank resource under an approved data request (ref: 32568). The authors declare no competing financial interests.

