## Supplemental Material for "The schizophrenia risk locus in SLC39A8 alters brain metal transport and plasma glycosylation"

**Supplementary Material**

**Table of Contents:**

| | Page # |
|---|---|
| Supp. Fig. 1. Use of MRI T2w/T1w data to classify by rs13107325 genotype. | 3 |
| Supp. Fig. 2. rs13107325 genotype has no effect on trace elements concentrations determined by ICP-MS except for Mn. | 4 |
| Supp. Fig. 3. Serum Mn concentrations are similar in males and females based on rs13107325 genotype. | 5 |
| Supp. Fig. 4. Serum Mn concentration does not correlate with age or BMI. | 6 |
| Supp. Fig. 5. Full MALDI-TOF spectra of N-glycans from severe SLC39A8 mutation carriers pre- and post- Mn supplementation. | 7 |
| Supp. Table 1. Clinical characteristics of Biobank participants based on rs13107325 genotype. | 8 |
| Supp. Table 2. Serum trace element ICP-MS Method Detection Limits. | 9 |

Supp. Table 3. Individual plasma protein N-glycan abundance based on rs13107325 genotype. 10

Supp. Table 4. Plasma protein N-glycan structure, name, mass, and characteristics used in study. 10

Supp. Table 5. Gender-based sub-analysis of branching plasma protein N-glycan abundance. 11

Supp. Table 6. Plasma protein N-glycan composition based on rs13107325 genotype. 12

Supp. Table 7. Individual plasma protein N-glycan abundance in severe SLC39A8 mutation carriers pre- and post-Mn supplementation. 13

Supp. Table 8. Plasma protein N-glycan composition following Mn supplementation in severe SLC39A8 mutation carriers. 14

**Supplementary Figures:**

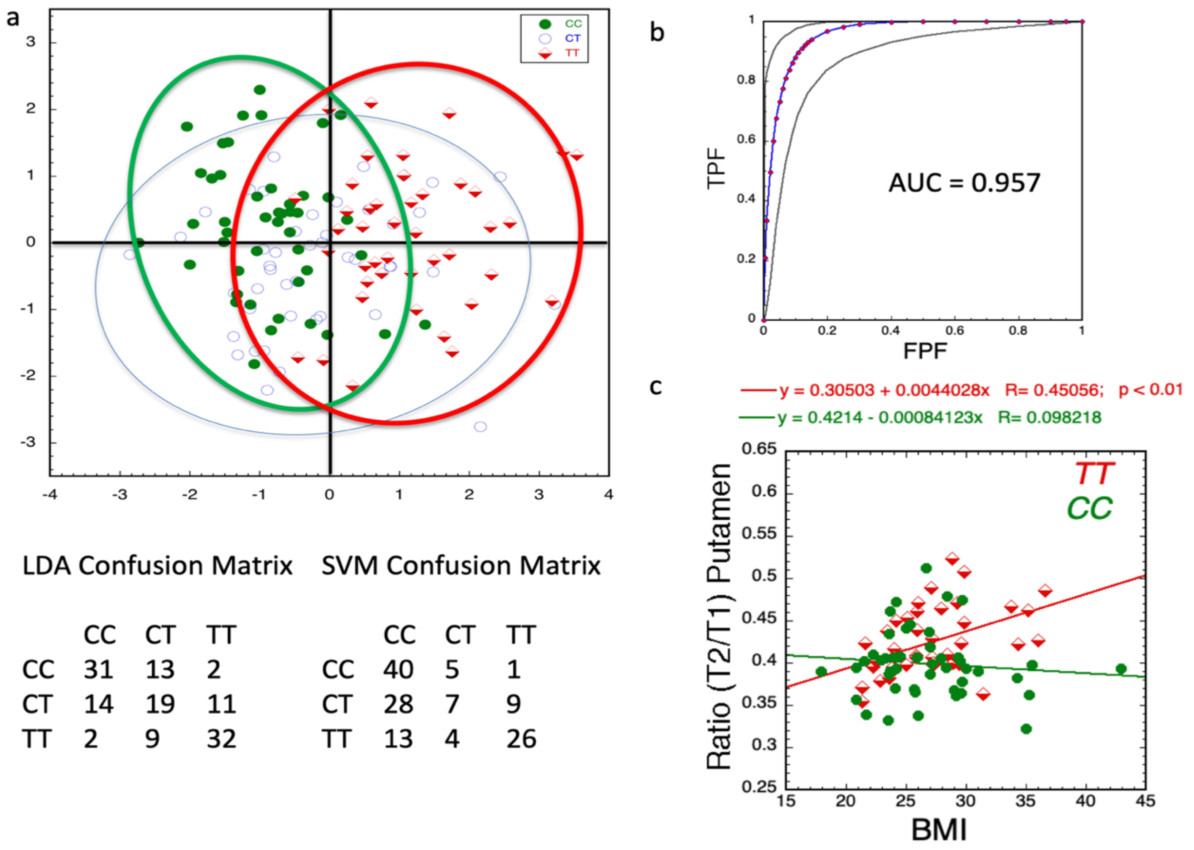


LDA Confusion Matrix

| | CC | CT | TT |
|---|---|---|---|
| CC | 31 | 13 | 2 |
| CT | 14 | 19 | 11 |
| TT | 2 | 9 | 32 |

SVM Confusion Matrix

| | CC | CT | TT |
|---|---|---|---|
| CC | 40 | 5 | 1 |
| CT | 28 | 7 | 9 |
| TT | 13 | 4 | 26 |

**Supplementary Figure 1**. **Use of MRI T2w/T1w data to classify by rs13107325 genotype.** A) Linear discriminant analysis (LDA) generated using the T2w/T1w ratio data in GPi, SN and LPut (three feature variables). There is excellent separation in along the primary axis between CC and TT genotypes, the CT carriers are spread out over the entire range (91% of the variance explained by axis one; separation along axis 2 was not significant by Wilk's lambda). We also performed support vector machines (SVM) classification with similar results as the LDA as seen in the confusion matrices using holdout analysis. B) ROC curve for binary classification between TT and CC carriers using the LDA classification. The area under the curve is 0.957 using only the three feature variables. C) We performed regression analyses using all the demographic data and the MRI data. The only significant effect was seen in the putamen which manifested a significant correlation with BMI (R = 0.45; p<0.01). The correlation between BMI and putamen for CC and CT carriers was not significant. CC n = 47, CT n = 45, TT n = 44.

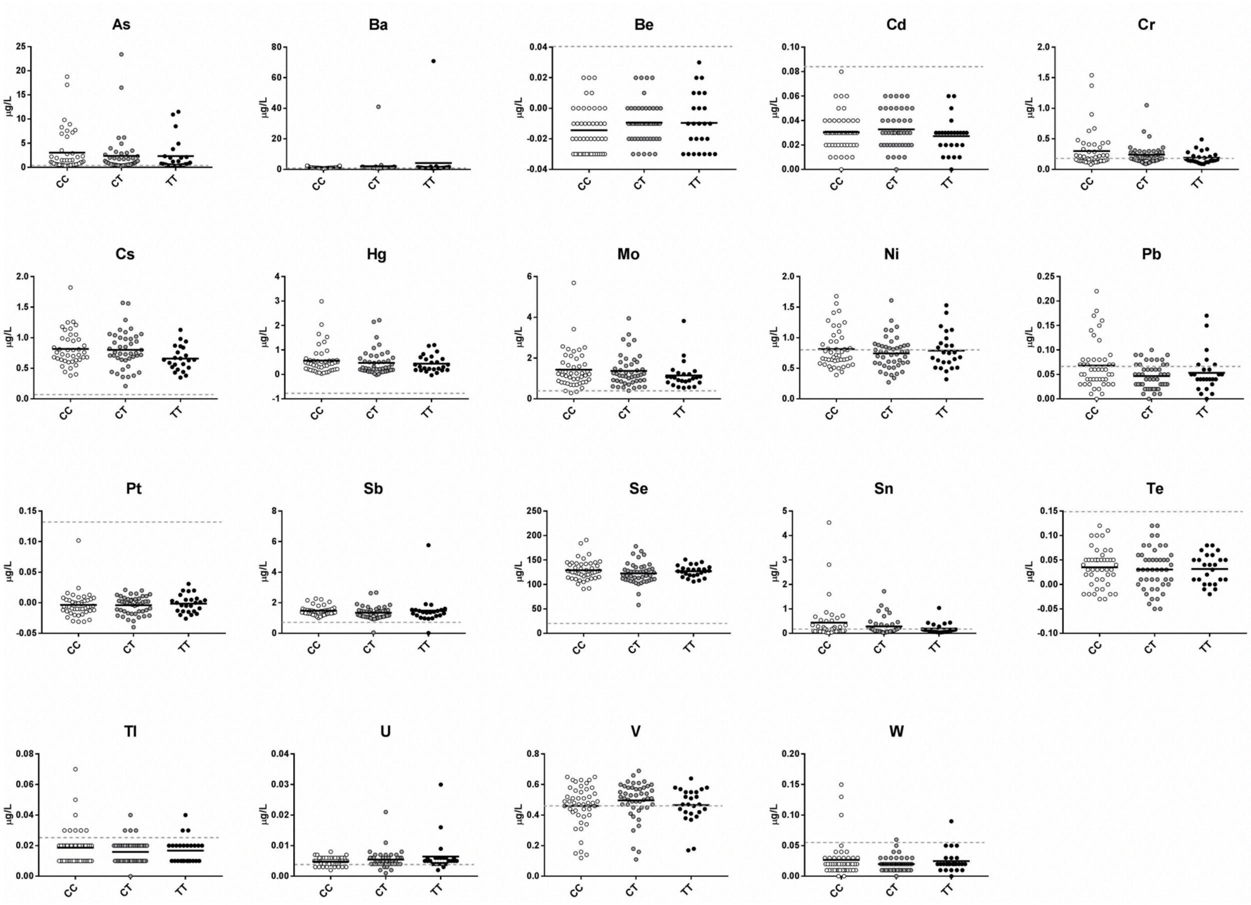


**Supplementary Figure 2**. **rs13107325 genotype has no effect on trace elements concentrations determined by ICP-MS except for Mn.** Data shown for each individual with black horizontal line representing mean for group. Method Detection Limit (MDL) shown as grey dashed line on each graph. (CC n = 46, CT n = 46, TT n = 25). *p-value <0.05, **p-value <0.01, ***p-value <0.001.

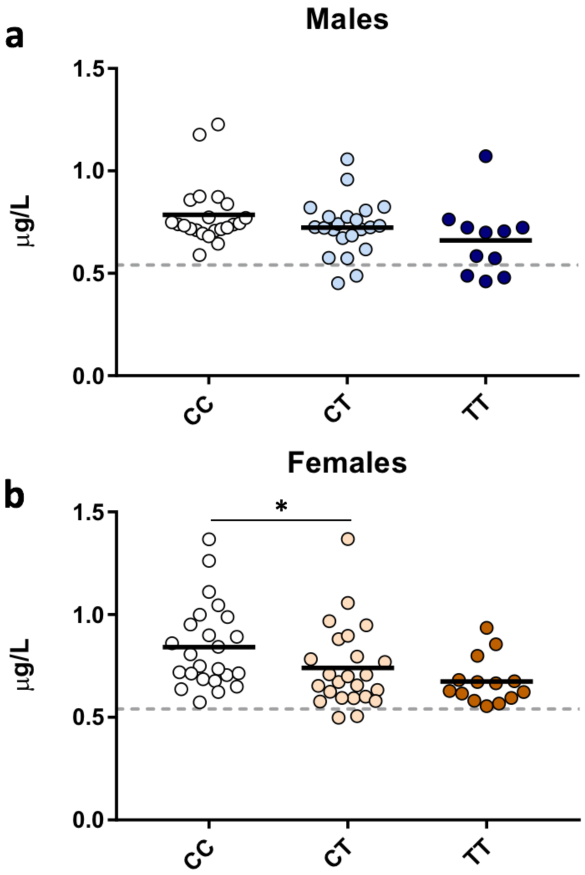


**Supplementary Figure 3**. **Serum Mn concentrations are similar in males and females based on rs13107325 genotype**. **a,** Males. **b,** Females. Data shown for each individual with black horizontal line representing mean for group. Method Detection Limit (MDL) shown as grey dashed line on each graph. (Males: CC n = 22, CT n = 22, TT n = 11) (Females: CC n = 24, CT n = 24, TT n = 14). *p-value <0.05, **p-value <0.01, ***p-value <0.001. Males: CC 0.785 μg/L vs CT 0.723 μg/L, p-value = 0.16, CC vs TT 0.661 μg/L, p-value = 0.06, CT vs TT p-value = 0.32); Females: CC 0.841 μg/L vs CT 0.740 μg/L, p-value = 0.08, CC vs TT 0.675 μg/L, p-value = 0.003, CT vs TT p-value = 0.20).

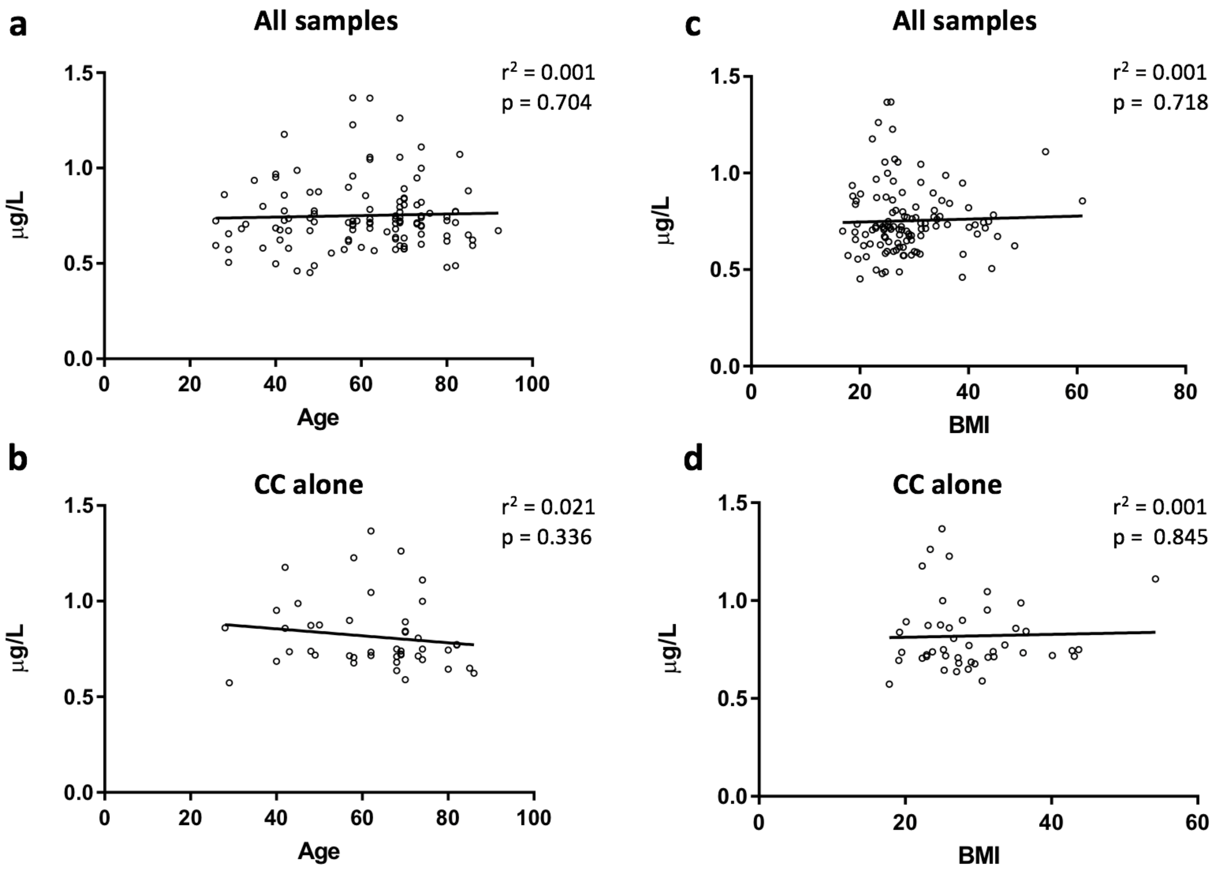


**Supplementary Figure 4**. **Serum Mn concentration does not correlate with age or BMI**. Linear regression of Mn concentration on age or BMI showed no significant correlation in (**a,c**) all samples or (**b,d**) CC genotype alone. Data shown for each individual result with black horizontal trend line. AGE: All samples n = 117, $r^2$ = 0.001, p = 0.704, Slope of best fit: 0.0004 +/- 0.0011, Slope 95% Confidence Interval: -0.0017 to 0.0025. CC Genotype n = 46, $r^2$ = 0.021, p = 0.336, Slope of best fit: -0.0018 +/- 0.0019, Slope 95% Confidence Interval: -0.0055 to 0.0020. BMI: All samples n = 116, $r^2$ = 0.001, p = 0.718, Slope of best fit: 0.0008 +/- 0.0021, Slope 95% Confidence Interval: -0.0034 to 0.0050. CC Genotype n = 46, $r^2$ = 0.0009, p = 0.845, Slope of best fit: 0.007 +/- 0.003, Slope 95% Confidence Interval: -0.0067 to 0.0082.

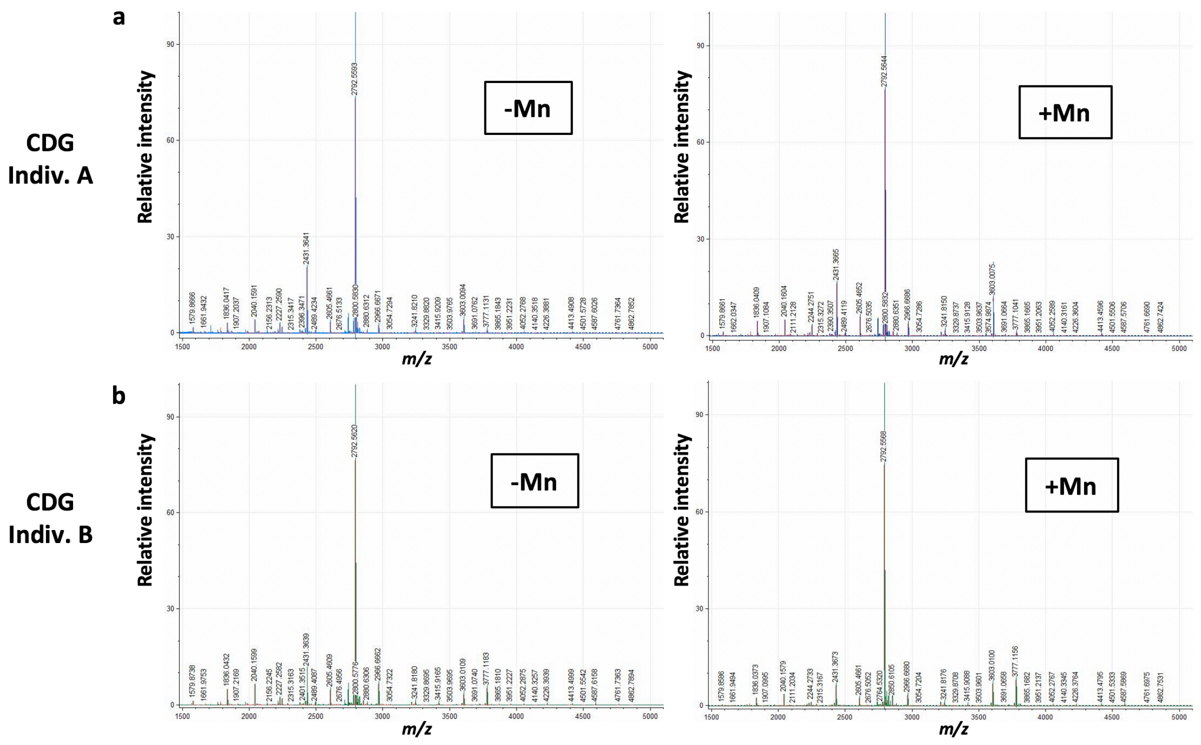


**Supplementary Figure 5. Full MALDI-TOF spectra of N-glycans from severe SLC39A8 mutation carriers pre- and post- Mn supplementation***.* MALDI-TOF spectrum from plasma/serum of (**a**) subject A and (**b**) subject B pre- and post-Mn supplementation. X-axis scaled for m/z of 1500-5000 kd and relative signal intensity on the Y-axis.

**Supplementary Tables:**

| UK BIOBANK | CC (n=46) | CT (n=48) | TT (n=48) |
|---|---|---|---|
| Age (SD) | 61.5 (6.8) | 61.4 (6.7) | 61.5 (6.9) |
| Gender (% Female) | 60% | 60% | 60% |
| BMI (SD) | 26.8 (4.6) | 27.1 (4.7) | 27.1 (4.8) |
| Smoker (% Yes) | 40% | 40% | 40% |
| Townsend Index (SD) | -2.1 (2.5) | -1.9 (2.5) | -2.1 (2.5) |
| **PARTNERS BIOBANK** | **CC (n=46)** | **CT (n=46)** | **TT (n=25)** |
| Age (SD) | 62.2 (14.5) | 62.2 (14.5) | 54.0 (18.6) |
| Gender (% Female) | 52% | 52% | 56% |
| BMI (SD) | 28.9 (7.5) | 29.2 (7.4) | 26.9 (6.26) |
| Smoker (% Yes) | 50% | 46% | 40% |

**Supplementary Table 1. Clinical comparisons between Biobank participants based on genotype.**

| Metal | µg/L |
|---|---|
| As | 0.24 |
| Ba | 0.21 |
| Be | 0.06 |
| Cd | 0.086 |
| Co | 0.313 |
| Cr | 0.18 |
| Cs | 0.048 |
| Cu | 110 |
| Hg | 0.14 |
| Mn | 0.54 |
| Mo | 0.32 |
| Ni | 0.76 |
| Pb | 0.06 |
| Pt | 0.22 |
| Sb | 0.049 |
| Se | 14 |
| Sn | 0.12 |
| Te | 0.15 |
| Tl | 0.025 |
| U | 0.004 |
| V | 0.46 |
| W | 0.053 |
| Zn | 42 |

**Supplementary Table 2. Serum trace element ICP-MS Method Detection Limits.** Based on 7 independent runs.

**Supplementary Table 3. Plasma protein N-glycan structure, name, mass, and characteristics used in study.**

PLEASE SEE ATTACHED EXCEL SHEET

**Supplementary Table 4. Individual plasma protein N-glycan abundance based on rs13107325 genotype.** #Heat maps scale: dark blue -> white -> bright red representing -50.0% -> 0 -> +50.0% for relative change.

PLEASE SEE ATTACHED EXCEL SHEET

| *ALL* | % Abundance | | | Change vs CC[A] | | % Change vs CC[B] | |
|---|---|---|---|---|---|---|---|
| **Antennarity** | **CC (n=33)** | **CT (n=31)** | **TT (n=25)** | **CT** | **TT** | **CT** | **TT** |
| **High-mannose** | 3.59 | 3.41 | 3.60 | -0.18 | 0.01 | -5 | 0 |
| **Mono-antennary** | 1.09 | 1.01 | 0.94 | -0.08 | -0.15 | -7 | -14 |
| **Bi-antennary** | 87.98 | 90.16** | 90.09** | 2.19 | 2.11 | 2 | 2 |
| **Tri-antennary** | 6.77 | 4.99* | 4.91* | -1.77 | -1.86 | -26 | -27 |
| **Tetra-antennary** | 0.57 | 0.42 | 0.46 | -0.15 | -0.12 | -27 | -20 |
| *MALE* | | | | | | | |
| **Antennarity** | **CC (n=17)** | **CT (n=15)** | **TT (n=10)** | **CT** | **TT** | **CC vs CT** | **CC vs TT** |
| **High-mannose** | 2.95 | 3.61 | 2.94 | 0.66 | -0.01 | 22 | 0 |
| **Mono-antennary** | 0.96 | 0.96 | 0.94 | 0.01 | -0.02 | 1 | -2 |
| **Bi-antennary** | 87.91 | 90.74* | 91.43 | 2.83 | 3.52 | 3 | 4 |
| **Tri-antennary** | 7.55 | 4.39** | 4.28** | -3.15 | -3.27 | -42 | -43 |
| **Tetra-antennary** | 0.64 | 0.30* | 0.42 | -0.34 | -0.22 | -53 | -34 |
| *FEMALE* | | | | | | | |
| **Antennarity** | **CC (n=16)** | **CT (n=16)** | **TT (n=12)** | **CT** | **TT** | **CC vs CT** | **CC vs TT** |
| **High-mannose** | 4.27 | 3.23 | 4.07 | -1.05 | -0.20 | -24 | -5 |
| **Mono-antennary** | 1.24 | 1.06 | 0.95 | -0.17 | -0.29 | -14 | -24 |
| **Bi-antennary** | 88.05 | 89.62 | 89.14 | 1.58 | 1.09 | 2 | 1 |
| **Tri-antennary** | 5.94 | 5.55 | 5.36 | -0.38 | -0.57 | -6 | -10 |
| **Tetra-antennary** | 0.50 | 0.53 | 0.48 | 0.03 | -0.02 | 5 | -4 |

**Supplementary Table 5. Gender-based sub-analysis of branching plasma protein N-glycan abundance.** [#]Heat maps scale: dark blue -> white -> bright red representing -5.0-> 0 -> +5.0 for absolute abundance change. *p-value <0.05, **p-value <0.01 for % abundance of CT and TT vs CC genotype.

| | % Abundance | | | Change vs CC[A] | | % Change vs CC[B] | |
|---|---|---|---|---|---|---|---|
| **Glycan Classes** | **CC (n=33)** | **CT (n=31)** | **TT (n=25)** | **CT** | **TT** | **CT** | **TT** |
| **Hybrid** | 0.76 | 0.74 | 0.69 | -0.02 | -0.07 | -2 | -9 |
| **Bisecting** | 7.83 | 7.05 | 7.84 | -0.78 | 0.01 | -10 | 0 |
| **Core Fucose** | 29.58 | 29.86 | 30.79 | 0.28 | 1.21 | 1 | 4 |
| **Antennary Fucose** | 2.09 | 1.56 | 1.43* | -0.54 | -0.67 | -26 | -32 |
| **-1 GlcNAc** | 1.42 | 1.42 | 1.34 | 0.00 | -0.08 | 0 | -5 |
| **Full Gal** | 81.04 | 80.30 | 81.12 | -0.73 | 0.08 | -1 | 0 |
| **-1 Gal** | 10.62 | 11.80 | 11.73 | 1.18 | 1.11 | 11 | 10 |
| **-2 Gal** | 8.34 | 7.89 | 7.15 | -0.45 | -1.19 | -5 | -14 |
| **Full NeuAc** | 66.18 | 62.79 | 63.65 | -3.39 | -2.53 | -5 | -4 |
| **-1 NeuAc** | 29.37 | 32.05 | 30.88 | 2.68 | 1.52 | 9 | 5 |
| **-2 NeuAc** | 4.39 | 5.10 | 5.41 | 0.71 | 1.02 | 16 | 23 |
| **Glycans containing:** | **CC (n=33)** | **CT (n=31)** | **TT (n=25)** | **CT** | **TT** | **CT** | **TT** |
| **2 GlcNAc** | 3.59 | 3.41 | 3.60 | -0.18 | 0.01 | -5 | 0 |
| **3 GlcNAc** | 0.89 | 0.82 | 0.75 | -0.07 | -0.14 | -8 | -16 |
| **4 GlcNAc** | 80.56 | 83.51* | 82.64 | 2.95 | 2.09 | 4 | 3 |
| **5 GlcNAc** | 14.37 | 11.83* | 12.54 | -2.54 | -1.84 | -18 | -13 |
| **6 GlcNAc** | 0.57 | 0.42 | 0.46 | -0.15 | -0.11 | -27 | -20 |
| **7 GlcNAc** | 0.01 | 0.01 | 0.01 | 0.00 | 0.00 | -33 | -13 |
| **0 Gal** | 12.02 | 11.38 | 10.83 | -0.64 | -1.19 | -5 | -10 |
| **1 Gal** | 11.55 | 12.66 | 12.52 | 1.11 | 0.97 | 10 | 8 |
| **2 Gal** | 69.09 | 70.55 | 71.28 | 1.45 | 2.19 | 2 | 3 |
| **3 Gal** | 6.77 | 4.99* | 4.91* | -1.77 | -1.86 | -26 | -27 |
| **4 Gal** | 0.56 | 0.41 | 0.45 | -0.15 | -0.12 | -27 | -21 |
| **5 Gal** | 0.012 | 0.008 | 0.010 | 0.00 | 0.00 | -33 | -13 |
| **0 NeuAc** | 25.25 | 26.44 | 26.13 | 1.18 | 0.88 | 5 | 3 |
| **1 NeuAc** | 21.27 | 22.96 | 21.93 | 1.69 | 0.66 | 8 | 3 |
| **2 NeuAc** | 47.79 | 46.80 | 48.01 | -1.00 | 0.22 | -2 | 0 |
| **3 NeuAc** | 5.50 | 3.71** | 3.79* | -1.79 | -1.71 | -33 | -31 |
| **4 NeuAc** | 0.18 | 0.09* | 0.13 | -0.09 | -0.05 | -51 | -27 |
| **0 Fuc** | 68.37 | 68.62 | 67.82 | 0.25 | -0.55 | 0 | -1 |
| **1 Fuc** | 31.58 | 31.34 | 32.15 | -0.24 | 0.57 | -1 | 2 |
| **2 Fuc** | 0.05 | 0.04 | 0.03 | -0.01 | -0.01 | -16 | -30 |
| **Monosacharride %** | **CC (n=33)** | **CT (n=31)** | **TT (n=25)** | **CT** | **TT** | **CT** | **TT** |
| **Man** | 30.46 | 30.62 | 30.56 | 0.16 | 0.10 | 1 | 0 |
| **GlcNAc** | 38.88 | 39.04 | 38.87 | 0.17 | 0.00 | 0 | 0 |
| **Gal** | 15.62 | 15.67 | 15.73 | 0.05 | 0.10 | 0 | 1 |
| **NeuAc** | 11.83 | 11.44 | 11.57 | -0.39 | -0.26 | -3 | -2 |
| **Fuc** | 3.21 | 3.23 | 3.27 | 0.02 | 0.06 | 1 | 2 |

**Supplementary Table 6. Plasma protein N-glycan composition based on rs13107325 genotype.**

[#]Heat maps scale: dark blue -> white -> bright red representing -5.0-> 0 -> +5.0 for absolute abundance change and -50.0% -> 0 -> +50.0% for relative change.

**Supplementary Table 7. Individual plasma protein N-glycan abundance in severe SLC39A8 mutation carriers pre- and post-Mn supplementation.** *Three masses in BOLD highlight glycans elevated in other MALDI-TOF studies of CDGs.

PLEASE SEE ATTACHED EXCEL SHEET

| | % Abundance | | | | % Change + Mn[A] | | Relative Change + Mn[B] | |
|---|---|---|---|---|---|---|---|---|
| Glycan Classes | A | A + Mn | B | B + Mn | A + Mn | B + Mn | A + Mn | B + Mn |
| Hybrid | 0.95 | 0.97 | 0.63 | 0.71 | 0.02 | 0.07 | 2 | 12 |
| Bisecting | 1.10 | 3.90 | 3.26 | 4.35 | 2.80 | 1.10 | 256 | 34 |
| Core Fucose | 12.48 | 19.51 | 21.30 | 14.80 | 7.03 | -6.50 | 56 | -31 |
| Antennary Fucose | 0.98 | 1.03 | 4.99 | 7.79 | 0.05 | 2.80 | 5 | 56 |
| -1 GlcNAc | 1.91 | 1.03 | 1.44 | 0.48 | -0.88 | -0.95 | -46 | -66 |
| Full Gal | 90.45 | 90.96 | 88.46 | 93.10 | 0.51 | 4.65 | 1 | 5 |
| -1 Gal | 6.52 | 5.27 | 7.60 | 4.53 | -1.25 | -3.08 | -19 | -40 |
| -2 Gal | 3.03 | 3.77 | 3.94 | 2.37 | 0.74 | -1.57 | 24 | -40 |
| Full NeuAc | 73.24 | 73.18 | 77.20 | 83.35 | -0.06 | 6.15 | 0 | 8 |
| -1 NeuAc | 23.73 | 23.06 | 20.14 | 14.55 | -0.67 | -5.58 | -3 | -28 |
| -2 NeuAc | 3.03 | 3.75 | 2.66 | 2.09 | 0.73 | -0.57 | 24 | -21 |
| Glycans containing: | A | A + Mn | B | B + Mn | A + Mn | B + Mn | A + Mn | B + Mn |
| 2 GlcNAc | 3.76 | 2.39 | 2.28 | 1.31 | -1.37 | -0.96 | -36 | -42 |
| 3 GlcNAc | 1.31 | 0.95 | 0.84 | 0.39 | -0.36 | -0.45 | -28 | -54 |
| 4 GlcNAc | 88.09 | 81.30 | 84.90 | 78.93 | -6.79 | -5.97 | -8 | -7 |
| 5 GlcNAc | 6.38 | 14.57 | 11.25 | 18.47 | 8.19 | 7.22 | 128 | 64 |
| 6 GlcNAc | 0.46 | 0.78 | 0.72 | 0.89 | 0.32 | 0.16 | 70 | 22 |
| 7 GlcNAc | 0.00 | 0.01 | 0.00 | 0.01 | 0.01 | 0.00 | 164 | 110 |
| 0 Gal | 6.85 | 6.21 | 6.29 | 3.73 | -0.64 | -2.56 | -9 | -41 |
| 1 Gal | 7.89 | 6.29 | 8.52 | 5.05 | -1.60 | -3.47 | -20 | -41 |
| 2 Gal | 79.32 | 75.87 | 76.25 | 75.99 | -3.45 | -0.26 | -4 | 0 |
| 3 Gal | 5.48 | 10.84 | 8.22 | 14.35 | 5.36 | 6.13 | 98 | 75 |
| 4 Gal | 0.45 | 0.77 | 0.72 | 0.88 | 0.32 | 0.16 | 70 | 22 |
| 5 Gal | 0.00 | 0.01 | 0.00 | 0.01 | 0.01 | 0.00 | 164 | 110 |
| 0 NeuAc | 12.99 | 13.34 | 13.59 | 8.58 | 0.35 | -5.01 | 3 | -37 |
| 1 NeuAc | 23.30 | 19.79 | 16.77 | 11.43 | -3.51 | -5.34 | -15 | -32 |
| 2 NeuAc | 59.51 | 57.55 | 62.72 | 66.60 | -1.96 | 3.88 | -3 | 6 |
| 3 NeuAc | 4.10 | 9.03 | 6.63 | 13.02 | 4.94 | 6.39 | 120 | 96 |
| 4 NeuAc | 0.11 | 0.29 | 0.29 | 0.37 | 0.18 | 0.08 | 170 | 29 |
| 0 Fuc | 86.55 | 79.47 | 73.88 | 77.57 | -7.09 | 3.69 | -8 | 5 |
| 1 Fuc | 13.43 | 20.53 | 25.96 | 22.28 | 7.09 | -3.68 | 53 | -14 |
| 2 Fuc | 0.01 | 0.01 | 0.16 | 0.15 | 0.00 | -0.01 | -42 | -4 |
| Monosacharride % | A | A + Mn | B | B + Mn | A + Mn | B + Mn | A + Mn | B + Mn |
| Man | 30.18 | 28.89 | 28.89 | 27.56 | -1.29 | -1.33 | -4 | -5 |
| GlcNAc | 37.58 | 37.62 | 37.54 | 37.02 | 0.05 | -0.51 | 0 | -1 |
| Gal | 16.92 | 17.21 | 16.79 | 17.61 | 0.29 | 0.82 | 2 | 5 |
| NeuAc | 13.97 | 14.26 | 14.32 | 15.85 | 0.28 | 1.52 | 2 | 11 |
| Fuc | 1.35 | 2.02 | 2.46 | 1.96 | 0.67 | -0.49 | 50 | -20 |

**Supplementary Table 8. Plasma protein N-glycan composition following Mn supplementation in severe SLC39A8 mutation carriers.** [#]Heat maps scale: dark blue -> white -> bright red representing -5.0-> 0 -> +5.0 for absolute abundance change and -50.0% -> 0 -> +50.0% for relative change.
